# Temperature modulates stress response in anammox reactors

**DOI:** 10.1101/2020.02.17.952358

**Authors:** Robert Niederdorfer, Damian Hausherr, Alejandro Palomo, Jing Wei, Paul Magyar, Barth F. Smets, Adriano Joss, Helmut Bürgmann

## Abstract

Autotrophic nitrogen removal by anaerobic ammonium oxidizing (anammox) bacteria is an energy-efficient nitrogen removal process in wastewater treatment. However, full-scale deployment under mainstream conditions remains challenging for practitioners due to the high stress susceptibility of anammox bacteria towards fluctuations in dissolved oxygen and temperature. Here, we investigated the response of microbial biofilms with verified anammox activity to oxygen shocks under favorable and cold temperature regimes. Genome-centric metagenomics and metatranscriptomics were used to investigate the stress response on various biological levels. We show that temperature regime and strength of oxygen perturbations induced divergent responses from the process level down to the transcriptional profile of individual taxa. Temperature induced distinct transcriptional states in compositionally identical communities and transient pulses of dissolved oxygen resulted in the upregulation of stress-response only under favorable temperatures. Anammox species and other key biofilm taxa display different transcriptional responses to the induced stress regimes.

## Introduction

Autotrophic nitrogen removal by biological anaerobic ammonium oxidation (anammox) is increasingly implemented as an energy-efficient mechanism of fixed nitrogen elimination during wastewater treatment. Deployed for mainstream wastewater treatment plants (WWTPs), it may even permit operation under energy autarky^1^. In contrast to conventional nitrification-denitrification system, external carbon sources are not required to reach very low effluent nitrogen concentration, less aeration is needed and most of the organic load can be diverted for valorization e.g. biogas or bio-plastics production^2–4^. Autotrophic N removal with anammox involves the simultaneous oxidation of ammonium (NH_4_^+^) and reduction of nitrite (NO_2_^-^) under oxygen-limiting conditions^5^. In engineered systems aerobic ammonia-oxidizing bacteria (AOB) oxidize a fraction of the available NH_4_^+^ to NO_2_^-^ (nitritation), which is subsequently used as a terminal electron acceptor by anammox bacteria to oxidize the remaining NH_4_^+^ to N_2_ ^6^. The process can be realized in single stage^7^ or two-stage bioreactor systems^8^. Currently, autotrophic N removal with anammox coupled to nitritation is already widely applied and represents a robust method for the treatment of wastewaters with high N concentrations under mesophilic conditions, e.g. effluents from anaerobic sludge digestion^9^. However, development of stable anammox processes for mainstream municipal WWTPs suffers from unexplained process instabilities due to unexpected fluctuation of environmental temperature and dissolved oxygen^10–12^.

Anammox bacteria are characterized by very slow growth rates, low cell yields^13^, and a high sensitivity to changing environmental conditions^14^. For their application in wastewater treatment, they are grown either in biofilm reactors on various carrier materials or as granules to retain sufficient biomass^15^. All respective reactor configurations support the formation of complex microbial communities^16^ with many potential synergistic and antagonistic interactions^17, 18^. Anammox, nitrification, heterotrophic denitrification, ammonification, as well as N incorporation may occur simultaneously within the system^19, 20^, thus complicating efforts to disentangle the sources of process instabilities.

Several studies have identified environmental stressors that affect the anammox process on the performance level (reviewed in Jin et al. 2012)^21^. Transient pulses of common wastewater constituents in the influent, such as heavy metals, phosphates, high NO_2_^-^ concentration, and sulfides have been reported as problematic factors, causing process instabilities^10, 11, 22^. However, dissolved oxygen (DO) and temperature were frequently discussed as the most critical factors with regards to stable operation of the anammox process^10, 23–26^. Low temperature is a seasonal stressor especially for mainstream anammox^27^. In contrast, even short term oxygenation of anammox systems have been associated with system failures ^26^. Oxygen control is of particular importance when it comes to the operation of the PN/A process in a single reactor. Numerous studies implementing intermittent aeration strategies have however demonstrated that successful nitritation and anammox processes are possible under these conditions^28, 29^.

So far, the majority of these studies were focusing on how disturbances affect the autotrophic N removal efficiency on the process level but did not investigate effects on microbial guild dynamics and their respective stress response, which might have led to the observed performance failures and/or recovery processes. Metagenomic shotgun sequencing allows to gain a comprehensive insight in the functional potential of communities involved in the anammox process^17, 18, 30, 31^ but is unlikely to yield useful information on short-term disturbance effects, which, in most of the cases, are not leading to shifts in the microbial community composition. Therefore we focus on biochemical measurements in combination with metatranscriptomics, which can provide important information on the response of microbes in a variable environment^32^. Understanding the transcriptional response of microbial communities but also its individual members to stress and how this response links to process-level effects is necessary for understanding process stability. A better understanding of the processes underlying system failure or stability are crucial for successful implementation of autotrophic N removal at full scale.

Here, we investigated the performance dynamics of complex anammox biofilms in response to short-term DO perturbations under two different temperature regimes. We hypothesized (1) that biofilms already exposed to certain stressors (e.g. lower temperatures) would experience stronger and longer lasting disturbance of the anammox process by the applied DO shocks. We further hypothesized (2) that DO shocks would result in a specific transcriptional response of the biofilm community in general, and anammox bacteria specifically. By analyzing the transcriptional response of different species, we aim to obtain information on the stress-level experienced by different microorganisms. To verify the reproducibility of observed effects, we ran parallel disturbance experiments in three laboratory-scale sequencing batch reactors (SBR) under comparable conditions. By combining high resolution monitoring of performance parameters with omics-based analysis of community composition, biodiversity and gene transcription, we aimed to determine transcriptional mechanisms connected to process disruptions. By mapping metatranscriptome data to metagenome-assembled genomes of anammox bacteria and other major biofilm members in a bioreactor system for autotrophic N removal, this study demonstrates that the impact of disturbances can be determined down to the level of transcriptional activity in individual microbial species using meta-omics tools.

## Methods

### Reactor Setup

Triplicate 12 L temperature-controlled bioreactors for autotrophic N removal from mainstream WWTPs by biological anammox were operated in parallel. Reactors were inoculated with 1 kg of biofilm carriers (∼12×12 mm; surface area 1.200-1800 m² m^-^³; FLUOPUR^®,^ Wabag, Switzerland) with verified anammox activity obtained from a pilot-scale (8 m^3^) mainstream anammox reactor, which is exposed to the seasonal temperature variations of the inflowing wastewater (∼25-17 °C) and successfully operated for over 2 years (∼100 g_NH4-N_ m^3^ d^-1^). A chemical oxygen demand-depleted effluent from another pilot scale (8 m^3^) mainstream high rate activated sludge reactor (Eawag Dübendorf, Switzerland) was employed as influent^11^. The reactors were operated in SBR mode with online monitoring and data logging of sensor data on concentrations of NH_4_^+^, nitrate (NO_3_^-^), pH, temperature, conductivity, and redox potential^10, 12, 33^. Each SBR cycle of approximately 6.5 h was controlled by an automated control sequence that consisted of five steps: (1) settling (30 min), (2) effluent discharge, (3) feeding (6 L of pre-treated wastewater and addition of NH_4_Cl solution to a final concentration of 30 mg_NH4-N_·L^-1^), (4) reaction phase (variable duration, until NH_4_^+^ concentration reached 5 mg_NH4-N_·L^-1^) and (5) mixing (1 h to draw down residual NO_2_^-^). During the reaction phase, NO_2_^-^ was frequently added in small doses to maintain a concentration of 1-2 mg N L^-1^ to allow continuous anammox activity. Anoxic conditions were maintained by continuously purging with a mixture gas of 95% N_2_ and 5% CO_2_.

### Pulse disturbance experiments under different temperature regimes

Short-term DO perturbation experiments were conducted under different temperature regimes (20 °C, 14 °C) to investigate process variability and to assess temperature-dependent stress responses of anammox consortia (Figure 1A). For each temperature regime, the operational period of the reactors was 7 days. After experiments at 20 °C were concluded, reactors were emptied, cleaned and refilled with fresh carriers for the experiments at 14 °C. Before filling of the reactors, biofilm carriers were taken from the pilot scale reactor (Figure 1: Inoculum 1/2). Reactors were first operated continuously without disturbance, i.e. baseline conditions, for 3 days to determine process stability and reproducibility in performance characteristics. A transient DO perturbation was then applied by raising DO to 0.3 mg L^-1^ for one hour during the reaction phase of one SBR cycle. This was followed by 36 hours of undisturbed operation to allow the system to fully recover. Reactors were then exposed to one hour of 1.0 mg L^-1^ of DO. Finally, reactors were operated for three additional SBR cycles of undisturbed operation. After both oxygen exposure periods, reactors were immediately flushed with additional N_2_-gas to restore anoxic conditions.

**Figure 1.**
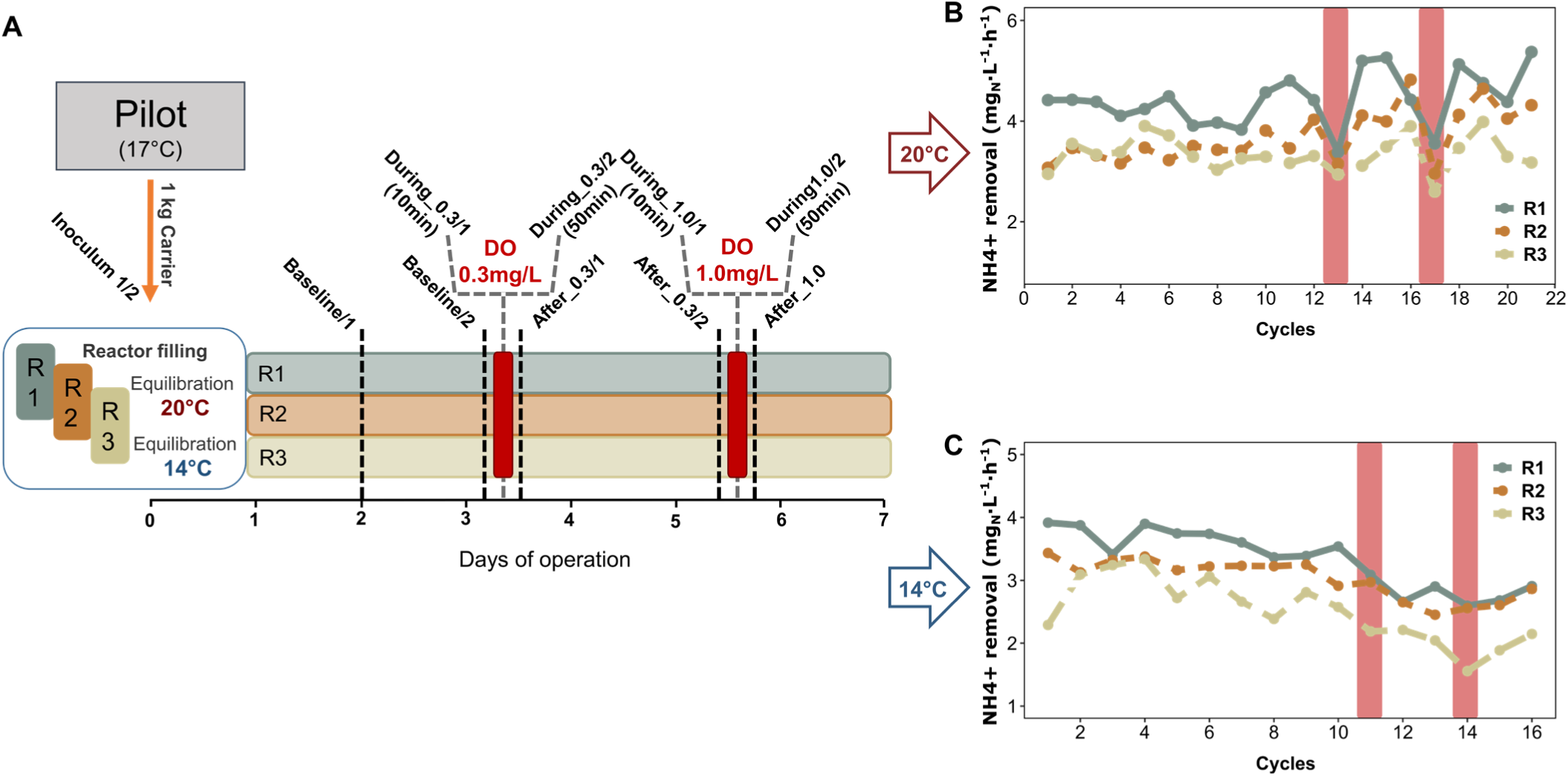
**A** Experimental set-up and carrier sampling scheme with sample names. **B** Average NH4+ removal per SBR cycle in mg Nitrogen/Liter/hour during the 20 °C experiment. Colors of the linecharts denote the different reactors (Green: Reactor1; Brown: Reactor2; Beige: Reactor3). Red bars highlight the DO disturbed cycles. **C** Average NH4^+^removal per SBR cycle mg Nitrogen/Liter/Hour during the 14 °C experiment. Colors of the lines denote the different reactors. Red bars highlight the DO disturbed cycles.

Anammox activity was defined as the volumetric NH_4_^+^ removal rate during the reaction phases of the SBR cycles. The NH_4_^+^ removal rate at each time point was calculated from the slope of a linear regression through the online concentration measurements by the NH_4_^+^ probes for a 10 min time interval.

The average SBR cycle duration during the experiments was 7.2±2.0 h at 20 °C and 8.8±2.3 h at 14 °C, respectively. The pH varied between 7.4 and 8.0 during the SBR cycles over the course of the experiments. Temperature remained stable in both experiments (avg. standard deviation: 0.25 °C (20 °C); 0.32 °C (14 °C)).

Samples for offline analysis were taken 4-5 times during a SBR cycle. Here, NH_4_^+^ concentrations were also determined with photochemical test kits (Hach Lange GmbH, Düsseldorf, Germany, Test LCK303, spectrophotometer type LASA 26) to recalibrate online NH_4_^+^ sensors if necessary. NO_2_^-^ concentrations were determined using colorimetric test strips (NO_2_^-^-test, 0-10 mg_NO2-N_·L^-1^, Merck KGaA, Darmstadt, Germany).

### Biomass sampling, extraction and sequencing

A sample of five biofilm carriers per reactor was taken on specific time points as shown in Figure 1A, resulting in a total number of 56 samples. Biofilm carriers were immediately snap frozen in liquid nitrogen and stored at −80 °C for later DNA and RNA extraction.

Nucleic acids were extracted based on a method modified from Griffiths et al.^34^. Biofilm carriers (n = 3) were cut into small pieces in a liquid nitrogen bath. Carrier pieces were transferred to 1.5 mL Matrix E lysis tubes (MPbio) and 0.5 mL of both hexadecyltrimethylammonium bromide buffer and phenol:chloroform:isoamylalcohol (25:24:1, pH 6.8) was added. Cells were lysed in a FastPrep machine (MPbio), followed by nucleic acid precipitation with PEG 6000 on ice. Nucleic acids were washed three times with ethanol (70%) and dissolved in 50 µL DEPC treated RNAse free water. Nucleic acids were separated overnight using a Lithium-Chloride (LiCl) solution (8 M). Resulting RNA pellets were purified and washed again three times with ethanol (70%) and dissolved in 50 µL DEPC treated RNAse free water. DNA was precipitated via Isopropanol from the LiCl supernatant. Pellets were washed two time with 70% ethanol and dissolved in 100 µL DEPC treated RNAse free water. DNA quality was assessed by using agarose gel electrophoresis and a Nanodrop ND-2000c (Thermo Fisher Scientific, USA). Total RNA quality and quantity was subsequently checked using the Agilent TapeStation system (Agilent, Santa Clara, CA, USA) to ensure only high-quality nucleic acids were used for downstream analysis. 100 ng RNA was used to construct strand-specific RNA-Seq libraries (Novogene, Hong Kong). 56 RNA samples (Sample IDs Figure 1) were sequenced on the Illumina HiSeq 4000 platform to generate 150 bp paired-end reads at greater sequencing depth. Nine DNA Samples (obtained at Baseline_2 (20 °C and 14 °C pooled for each reactor) and for both temperatures and each reactor at After_1.0) were sequenced on the Illumina NextSeq platform (Illumina, CA, USA) to generate 150 bp paired-end reads (350 bp mean insert size). All DNA and RNA sequencing was performed at Novogene, Hong Kong. Raw DNA and RNA sequences can be found on the MgRast server^35^ under project ID mgp87520.

### Metagenome assembly and annotation

Raw DNA sequencing reads were quality controlled with FastQC^36^ and Illumina adapters were trimmed, if necessary, with Trimmomatic^37^. Kaiju^38^ was used to taxonomically assign the raw reads, using maximum exact matches of the query sequences translated to amino acids and protein database sequences. The reads were aligned against the NCBI non-redundant (NR) protein sequences from all bacteria, archaea, viruses, fungi, and microscopic-sized eukaryotes, respectively. Paired-end reads from each sample were assembled with Megahit^39^ (∼550000 contigs; ∼N50: 1750 bp) and reads were mapped back via BBmap v35.92^40^ with the parameters minid = 0.95 and ambig = random to asses assembly quality and coverage. Resulting bam files were handled and converted as needed using SAMtools1.3^41^. Open reading frame (ORF) detection and subsequent gene annotation from assembled contigs was performed with prokka^42^. ORFs were queried against the SEED subsystems (pubseed.theseed.org; accessed July 2019), Clusters of Orthologous Groups (COG; https://www.ncbi.nlm.nih.gov/COG/; accessed June 2019) and Metacyc (https://metacyc.org/; accessed June 2019). To quantify the abundance of classified genes in the community. We mapped raw metagenomics reads back to the predicted and annotated ORFs of the assembled contigs. For comparability, we normalized the counts to genes per million. Here, counts were first normalized for gene length and afterward for sequencing depth. To ensure higher coverage for further transcriptomic mapping, we co-assembled the nine metagenomic samples.

### Recovery and assessment of metagenome-assembled genomes (MAGs)

High-quality trimmed reads from each sample were co-assembled into scaffolds using IDBA-UD^43^ with the options --pre_correction --min_contig 1000. metaWRAP^44^ binning and refinement modules were applied to the co-assembly. Completeness and contamination rates of the final bins were assessed using CheckM^45^. Bins were taxonomically classified using the metaWRAP classify module. Bin abundances were assessed using the metaWRAP quantification module. Here, raw reads were mapped against the putative genomes and abundance is expressed as the coverage of raw reads on the MAG. Phylogenetic analysis of the recovered draft genomes was performed with Phylosift v1.0.1, based on a set of 37 universal single-copy marker genes^46^ using the ‘phylosift -all’ option. Thirty publicly available genomes closely related to the recovered draft genomes and ecosystem were downloaded to build a phylogenetic tree. Concatenated amino acid sequences of the marker genes were aligned with Phylosift, and a maximum likelihood phylogenetic tree was constructed with RAxML v8.2.4 with the PROTGAMMAAUTO model and 100 bootstraps^47^. Metagenome assembled genomes can be accessed on European Nucleotide Archive (ENA) under accession no. PRJEB36638.

### Metatranscriptome analysis

Analysis of metatranscriptomic reads was performed according to Lawson et al. (2017)^17^. Quality filtered paired end reads were merged and rRNA sequences were filtered from the merged reads using SortMeRNA^48^ v2.0, based on multiple bacterial, archaeal and eukaryotic rRNA databases. Non-rRNA reads were mapped against the co-assembled metagenomic contigs (n = 9) using BBMap v35.92 with the parameters ‘minid = 0.95’, which specifies a minimum alignment identify of 95%. Ambiguous reads with multiple top-hit mapping locations were assigned to a random ORF (‘ambig = random’ option). Read counts were calculated for each predicted ORF using the FeatureCount option in the subread package^49^. Raw counts were normalized for sequencing depth and gene length and expressed as transcripts per million (TPM) ^50^ as a proxy for gene expression. The same workflow was applied to all recovered draft genomes to compare gene expression patterns across recovered MAGs. The relative transcript abundance (*a*) in TPM of the MAG was calculated according to Eq. 1 from the absolute number of cDNA reads that calculated from the total number of mRNA reads that mapped to the MAG, divided by the genome length (bp) and normalized for sequencing depth (TPM).

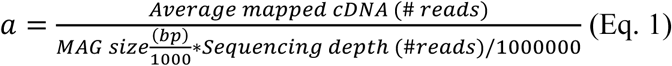

Furthermore, we investigated the relationship between MAG abundance and potential activity by calculating their ratio.

### Additional data analysis and statistics

Data for the replicate reactors are given as averages and standard deviation of all values. Normalized gene abundance and transcriptomic abundance tables from metagenome samples, draft genomes and metatranscriptomes, respectively, were used for all further bioinformatics and statistical analysis. All data analyses and visualization were conducted using R (R Core Team 2015, version 3.6)^51^ with the packages vegan (version 2.5-6)^52^ and ggplot2 (version 3.2.1)^53^.

MAG-level transcription was compared using the functional transcript abundance at the level of COG classes with a non-metric multi dimensional scaling analysis (nMDS). Here, we summed up all transcript abundances (TPM) corresponding to a COG category for each MAG. The obtained COG abundance matrix was used as input for the metamds (with the Jaccard option) function of the vegan package.

## Results

### Process level performance and disturbance response

Per-cycle reactor performance was comparable in all three reactors during the 3 days of operation under baseline conditions under both temperature regimes (Figure 1B and C). We did not observe anomalies in reactor activity potentially imposed by the carrier transfer from the pilot reactor (17 °C) to the small volume bioreactors (14 °C or 20 °C). The average NH_4_^+^ removal rates per hour within a SBR cycle during baseline conditions were 3.7±0.5 mg_NH4-N_·L^-1^ at 20 °C and 3.2±0.4 mg_NH4-N_·L^-1^ at 14 °C, respectively. However, the NH_4_^+^ removal rates of R1 (Figure 1B and C) were significantly higher compared to the other reactors independent of the temperature regime (p < 0.05, Student’s t-test). NH_4_^+^ removal rates in all reactors experienced similar trends over the course of the experiment. While at 20 °C the average NH_4_^+^ removal per hour within a cycle tended to increase after the baseline period (3.9±0.7 mg_NH4-N_·L^-1^, not significant), the reactor performance showed an opposite, decreasing trend at 14 °C to an average of 2.5±0.4 mg_NH4-N_·L^-1^ which was significant (linear regression, p < 0.005 for all reactors; R1: r^2^ = 0.8; R2: r^2^ = 0.7; R3: r^2^ = 0.6) (Figure 1B and C).

The DO perturbations caused an obvious impact on the average per cycle NH_4_^+^ removal rate at 20 °C, while the DO shocks at 14 °C did not impact the per cycle activity (Figure 1B and C red bars).

To understand the susceptibility of reactor performances to DO perturbations in more detail, we analyzed the SBR cycles and corresponding average NH_4_^+^ removal rates at a half-hour temporal resolution (Figure 2A and B). Here, a clear distinction in performance decay between high and low oxygen stress was observed. Pulse disturbances of 0.3 mg L^-1^ DO prompted only moderate declines in the average NH_4_^+^ removal rates (∼ 40% maximum activity loss) under both temperature regimes. On the other hand, 1.0 mg L^-1^ of DO completely inhibited NH_4_^+^ removal activity in all reactors at 20 °C (Figure 2A) but only partly at 14 °C (Figure 2B). However, after the 1.0 mg L^-1^ DO perturbation, all disturbance effects were rapidly reversible, even within the perturbed SBR cycle, except for R3 during the 14 °C experiment. The observed recover time to baseline performance averaged 34.7±4 min at 20 °C and 40.6±8 min at 14 °C, respectively (Supplementary Figure 1A). Interestingly, the time to reach maximum inhibition of the NH_4_^+^ removal rate was nearly the same for both temperature regimes and oxygen concentrations (Supplementary Figure 1B). On average, it took 72±4 min to reach the maximum inhibition of the NH_4_^+^ removal rate after start of the DO perturbation.

**Figure 2.**
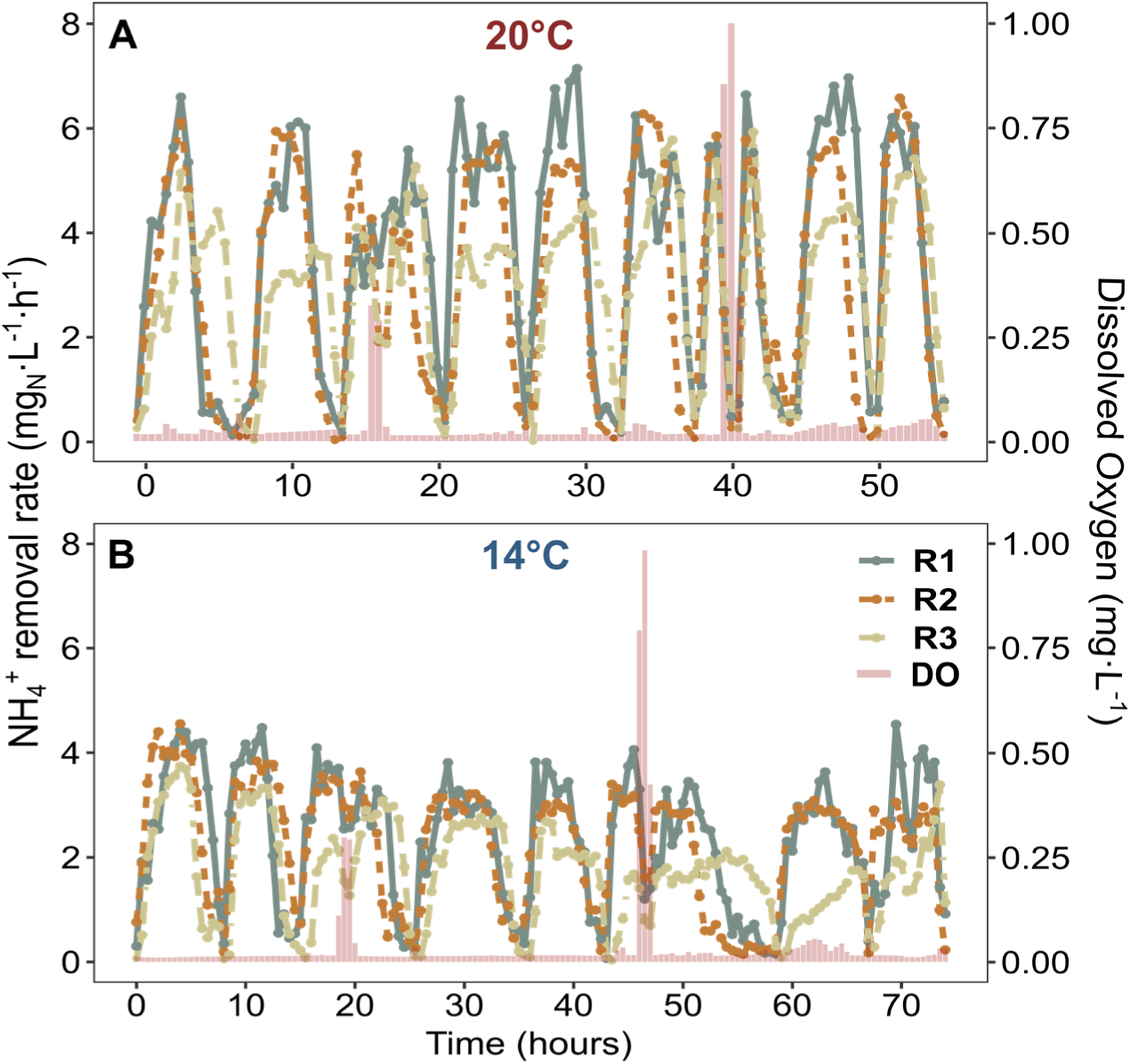
**A** NH_4_^+^ removal rates per hour over nine SBR cycles (20 °C). X-axis denotes the time in hours, the y-axis (left) displays the NH_4_^+^ removal rate in mg Nitrogen/Liter/hour. Colors of the linecharts denote the different reactors (Green: Reactor1; Brown: Reactor2; Beige: Reactor3). Concentration of dissolved Oxgen (red bars) in mg DO/Liter are shown on the 2nd y-axis (right). **B** NH_4_^+^ removal rates per hour over nine SBR cycles (14 °C).

### Microbial community structure and genetic potential

Based on the short-term nature of the experiment and generally weak disturbance effects we did not expect changes of the microbial community composition. Our metagenome analysis confirmed that microbial community composition and functional potential was stable (Figure 3A, Supplementary Figures 2, 3) as was the distribution of anammox species (Figure 3B). Differences between the samples, mainly on the species level are thought to primarily reflect carrier-to-carrier variation^54^.

**Figure 3.**
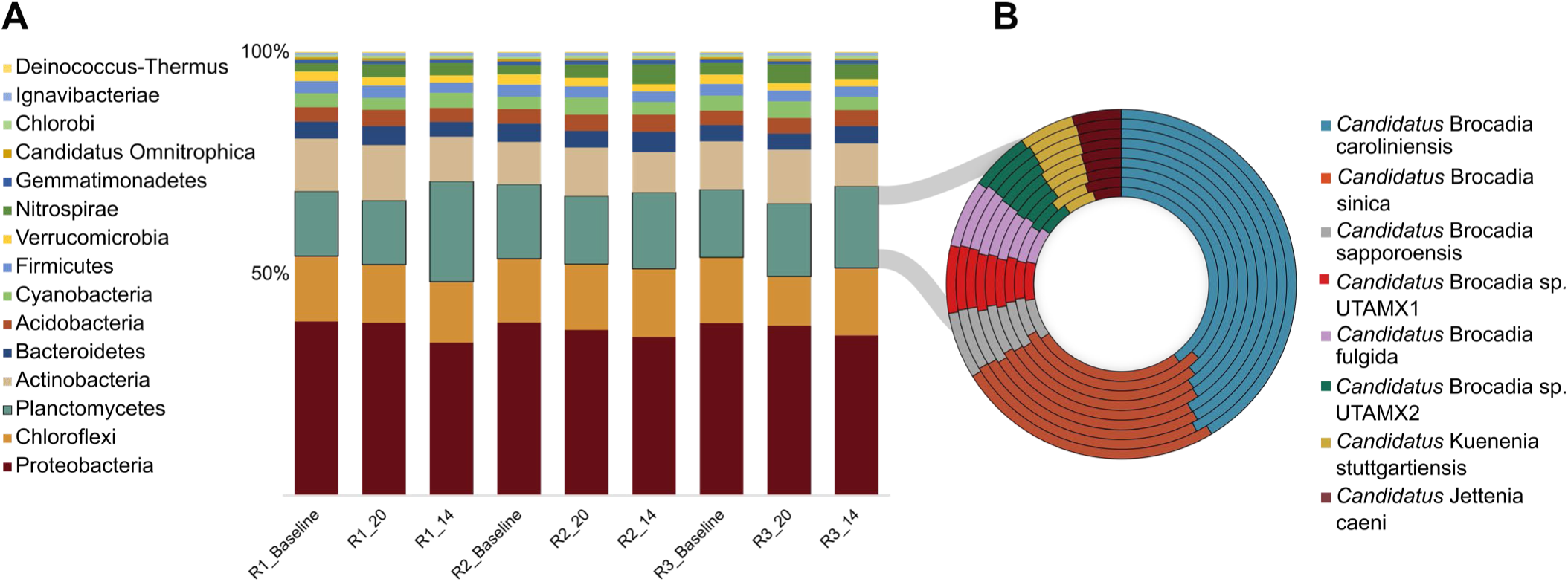
**A** Relative abundance of the top 15 bacterial phyla of the biofilm carrier community derived from metagenomic sequencing. Labels denote Reactor (R1, R2, R3) and time of sampling, _**Baseline** (Pooled from 20 °C and 14 °C experiment), **_20** (after the 20 °C experiment), **_14** (after the 14 °C experiment). Colors represent the different bacterial phyla. **B** Relative abundance of Anammox species within the samples derived from the same dataset – these species are the main contributors to the Planctomycetes phylum in (A). One ring is one samples and from the inner ring to outer ring samples follow the same order as in A (R1_baseline to R3_14 (n=9)).

The microbial community in the biofilm carriers consisted of a diverse assemblage of taxa, many of which are typical for mainstream anammox WWTP^17, 18, 30^. Members of Proteobacteria, Chloroflexi, Planctomycetes and Actinobacteria dominated the biofilms, accounting for approximately 70% of the total community composition, However, Proteobacteria represented the largest Phylum of the community accounting for ∼40% of the total abundance (Figure 3A). Other organisms that displayed moderate abundance were affiliated with Nitrospirae, Gemmatimonadetes and Acidobacteria.

Genera with known importance to the N cycle, e.g. anammox bacteria, denitrifying bacteria and nitrite oxidizing bacteria (NOB) were abundant with considerable internal diversity. In particular, the anammox species cluster was unexpectedly diverse with eight different anammox species affiliated to genera of *Candidatus* Brocadia, *Candidatus* Kuenenia and *Candidatus* Jettenia (Figure 3B). *Candidatus* Brocadia was the most abundant genus of the community (Supplementary Figure 2). Interestingly, NOB from the Genus *Nitrospira* accounted for ∼3% of the total community. Only a very small fraction of the community (∼0.4%) could be assigned to genera of aerobic AOB.

As expected, characterization of functional capacity of the community from contig gene content revealed that genes involved in COG energy production and conversion (C), protein synthesis (J, K, L), amino acid metabolism (E) and signal transduction (T) were the most abundant in the system (Supplementary Figure 3). For specific functional subsystems, we found that genes involved in the carbohydrate metabolism and production of vitamins and cofactors comprised the most abundant fractions. Genes involved in N transformations did not occur in high abundance (Supplementary Figure 4). However, genes belonging to the anammox metabolism (reddish colours Supplementary Figure 5) such as hydrazine dehydrogenase (*hdh*) and the hydrazine synthase cluster (*hzsαβγ*) and denitrification (blueish colours) were the most abundant ones within the N cycle category. To our surprise, also genes involved in dissimilatory nitrate reduction to ammonium (DNRA) occurred in moderate abundance, while assimilatory NO_3_^-^ reduction genes were only present in small numbers (Supplementary Figure 5).

Since no major shifts were observed in the community level nor in the functional potential between the nine metagenomic samples, a co-assembly was applied to ensure higher coverage for further transcriptomic analysis.

### Identity and abundance of Metagenome Assembled Genomes

To be able to dissect the ecology and stress response of individual bacterial lineages from the complex microbial communities used in these experiments the recovery of Metagenome Assembled Genomes (MAGs) is a necessary prerequisite. We investigated 12 MAGs ranging from medium to high quality (> 80% completeness, < 10% contamination) from the co-assembled metagenomic libraries. These MAGs were affiliated with the phyla Planctomycetes (AMX), Nitrospirae (NOB), Chloroflexi (CFX), Proteobacteria (APT), Bacteroidetes (BCD) and Gemmatimonadetes (GMT) (Table 1).

**Table 1.**
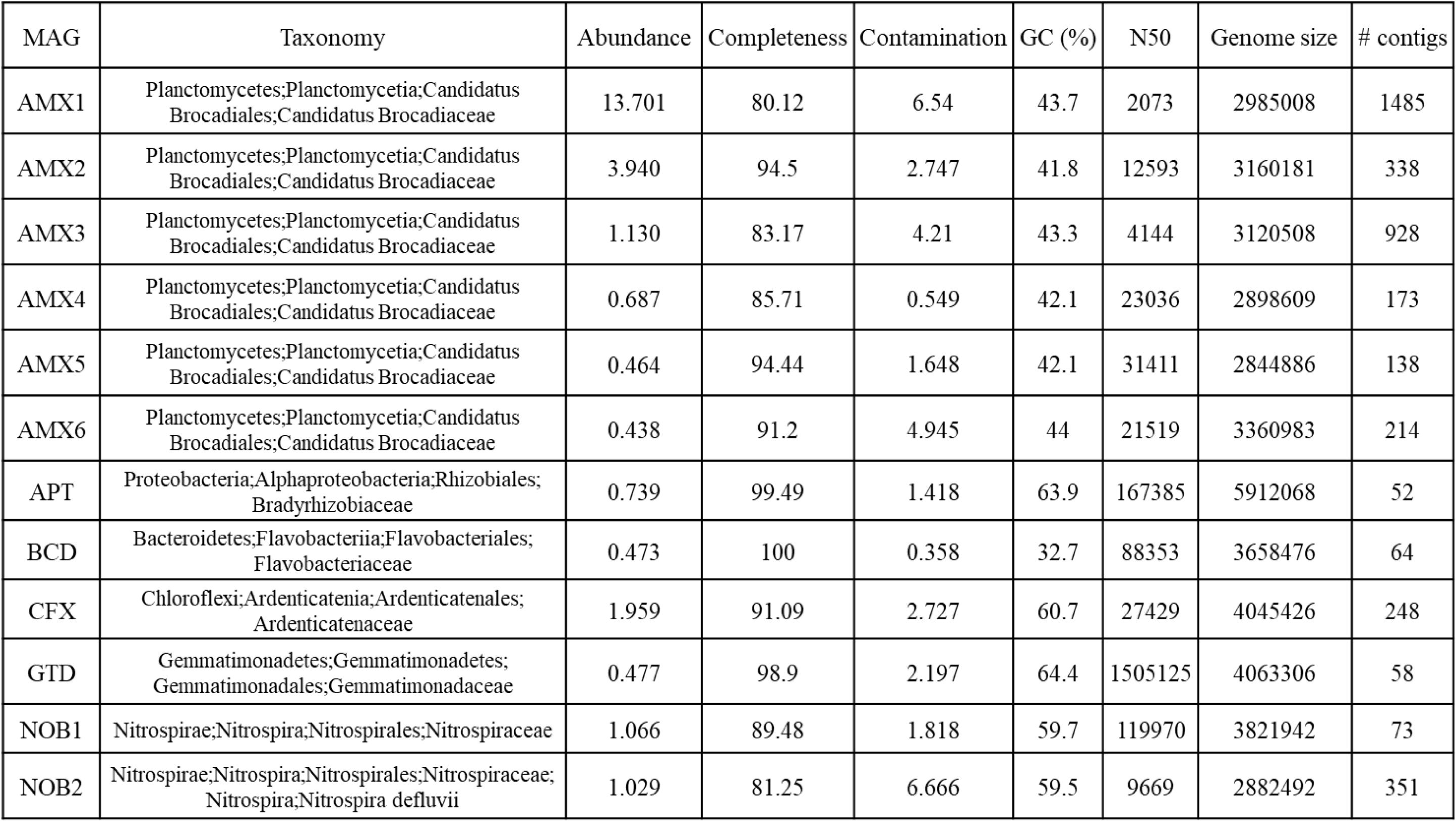
Binning statistics of MAGs recovered from the anammox community

The average relative abundance of the MAGs was calculated from the amount of total filtered DNA reads from each of nine metagenome samples that mapped to all retrieved MAGs. On average, the two most abundant MAGs accounted for 13.9% and 4%, of the community respectively (Table 1). The other MAG’s abundances ranged between 2.0 – 0.5%.

A phylogenetic tree based on the protein sequences of 37 single-copy marker genes^46^ (Supplementary Figure 6) showed that the majority of retrieved MAGs displayed high similarities to reference genomes from previous bioreactor studies^17, 18, 30^. Our results suggest that the six recovered putative anammox genomes (AMX1-6) are all representatives of the *Brocadia* genus and both NOB MAGs were closely related to *Nitrospira defluvii*. AMX4 and AMX2 created their own branch within the Genus *Brocadia*. The MAG affiliated with phylum Chloroflexi was assigned to the genus *Candidatus* Promineofilum, a well-known filamentous member of activated sludge communities with a facultative anaerobic lifestyle in WWTP^55^.

The taxonomic classification and relative abundance is also in line with our findings on the metagenome level, where anammox bacteria accounted for ∼20% of the community (Figure 3).

### MAGs and the N-cycle

The recovered MAGs were analysed with regard to their functional role in the bioreactors biological N-cycle network. As expected, all anammox MAGs carried at least two of the most important anammox genes, *hdh* and hydroxylamine oxidoreductase (*hao*) (Supplementary Table 1). However, *hzs* could not be annotated in AMX4 and AMX5. Interestingly, except for AMX3, none of the anammox MAGs encoded for *nirK* or *nirS* genes, which are typically responsible for the first step of the anammox process ^56^. The majority of retrieved anammox MAGs also contained *nrfH* genes (Supplementary Table 1), which encodes for enzymes responsible for the reduction of NO_2_^-^ to NH_4_^+^. We were not able to annotate the Nitrite:Nitrate Oxidoreductase (*nxr*) in the anammox MAGs likely due to its high sequence similarity to bacterial nitrate reductases^57^.

All other MAGs encoded capabilities for partial or full denitrification, DNRA and assimilatory NO_3_^-^ reduction (Supplementary Table 1). The first step in denitrification appears to be orchestrated by GTD and BCD carry the genes for respiratory nitrate reductase (*narGHI*) that reduces NO_3_^-^ to NO_2_^-^. The ability to reduce NO_2_^-^ to nitric oxide (NO) (*nirS*, *nirK*) was found in the Nitrospira (NOB1, NOB2), Chloroflexi and the Proteobacterium (APT) genomes. All of them contain either one or both variants of the nitrite reductases. BCD and all anammox genomes have the functional competence to reduce NO to nitrous oxide (N_2_O) via the nitric oxide reductase (*norB*, *norC*) to conclude the next step of partial denitrification. Finally, CFX, APT, GTD and BCD expressed genes for the reduction of N_2_O to N_2_ via nitrous-oxide reductase (*nosZ*). Thus, all retrieved MAGs seem to metabolically participate in the N cycle and are important for the overall N removal this ecosystem.

### Community-wide transcriptomic analysis

Metatranscriptomics allowed us to assess impacts of temperature regime and disturbance events on the community-wide transcriptional activity. RNAseq yielded on average 2 × 10^7^ reads per sample after quality filtering. Reads were mapped against the metagenomic co-assembly derived from the nine metagenomic samples to provide insight into the transcriptional response of the whole community.

The community-wide transcription differed significantly (PERMANOVA: p < 0.005) between the two temperature regime in all reactors. The nMDS ordination based on global transcript abundances (Figure 4) showed clustering by temperature, indicating a changed transcriptional status of the community. Further, all samples taken during experiments are clearly separated from the inoculum samples (17 °C pilot reactor). The dissimilarities between the two inoculum samples (yellow, orange), taken one week apart at the exact same time during a SBR cycle, indicate that variability in transcriptomic data has to be expected even without experimental intervention – either due to the dynamic nature of this engineered ecosystem or due to methodological error. These differences had no effect on the process level, as we did not observe significant differences in NH_4_^+^ removal rates between the two fillings of the experimental reactors.

**Figure 4.**
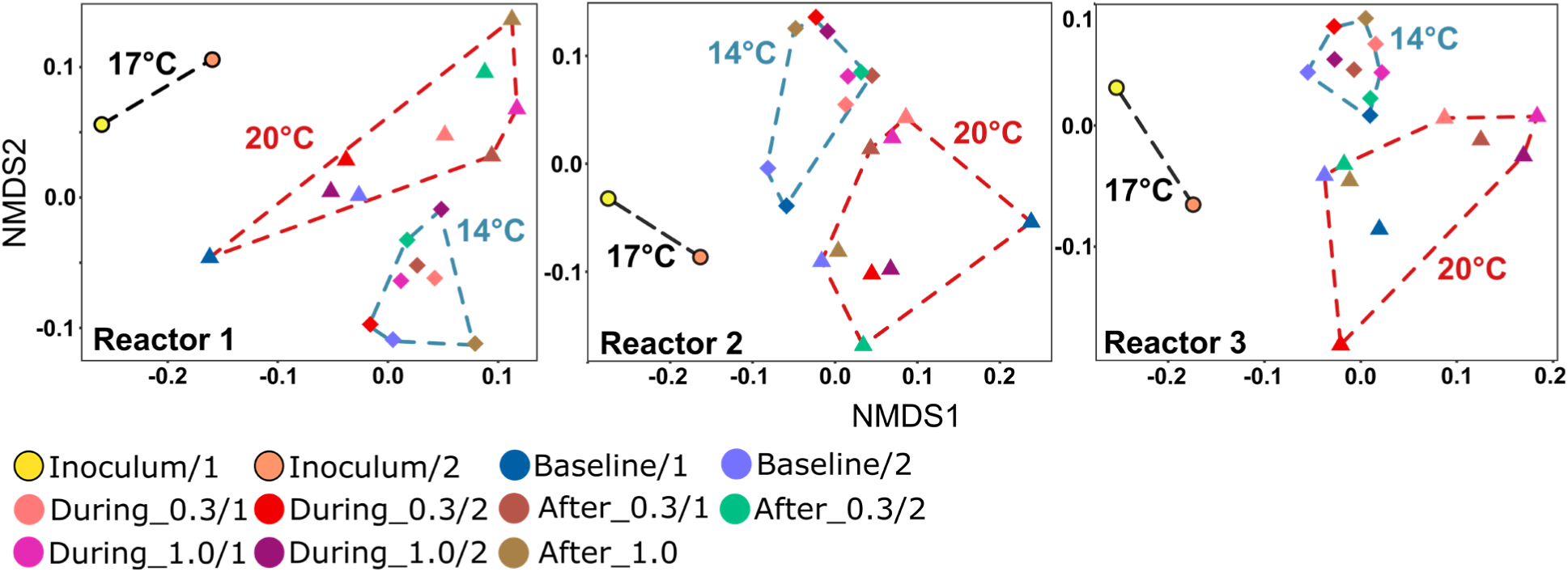
nMDS based on Jaccard dissimilarities depicting overall gene expression during the time-points of the experiment under different temperatures for all reactors. Colors of clusters denote the respective temperature of the experiment (black: Inoculum, red:20 °C, blue:14 °C). Reactor1 is missing one Baseline sample of the 14 °C experiment due to insufficient coverage of the metatranscriptome. Stress values: R1:0.12; R2:0.13; R3:0.12

DO disturbances did not induce a clear community-wide transcriptional response at either temperature regime, i.e. samples obtained at “stress” conditions did not cluster consistently apart from baseline samples (Figure 4). However, the transcriptional variance was consistently greater during the 20 °C experiment (greater hull area, Figure 4), than during the 14 °C experiment. Even for baseline samples, which were taken at the same stage in the SBR cycle one day apart, we found large variation in transcription under the 20 °C regime. Differences in transcriptional status between baseline conditions were less pronounced at 14 °C.

### Transcriptional response of functional gene categories

To explore further implications of temperature on transcript abundance and to assess if oxygen affected transcription of specific gene networks we investigated the influence of disturbances on gene transcription on the level of functional categories and for N metabolism, respectively.

Transcript abundance was, unsurprisingly, the highest for genes in the COG categories C (Energy production and conversion), E (Amino acid transport and metabolism), J (Translation), K (Transcription) and O (posttranslational modification) independent of the temperature regime (Figure 5A). In contrast to our findings on the global transcriptomic profile, DO disturbances during the 20 °C regime appeared to have notable impacts on some COG categories. Especially, genes within the clusters J, K and O displayed elevated relative transcription during the first 10 minutes of 0.3 mg L^-1^ DO (15%±3 increase), after the 0.3 mg L^-1^ DO disturbance (15±6% increase) and during the whole period of the 1.0 mg L^-1^ DO perturbation (26±7% and 26±4% increase, respectively). The most prominent genes within these categories were annotated as cold shock-like proteins (K) and a variety of ribosomal proteins, RNA polymerase sigma factors (J) and Heat-shock proteins (O). These genes displayed high fluctuations in relative transcription at 20 °C but maintained a stable transcription over the whole experiment at 14 °C. In line with findings for global gene transcription (Figure 4), changes in transcription of COG classes were in general much less pronounced at 14 °C. However, transcript abundances during baseline conditions were comparable independent of the temperature (p > 0.5, Student’s t-test).

**Figure 5.**
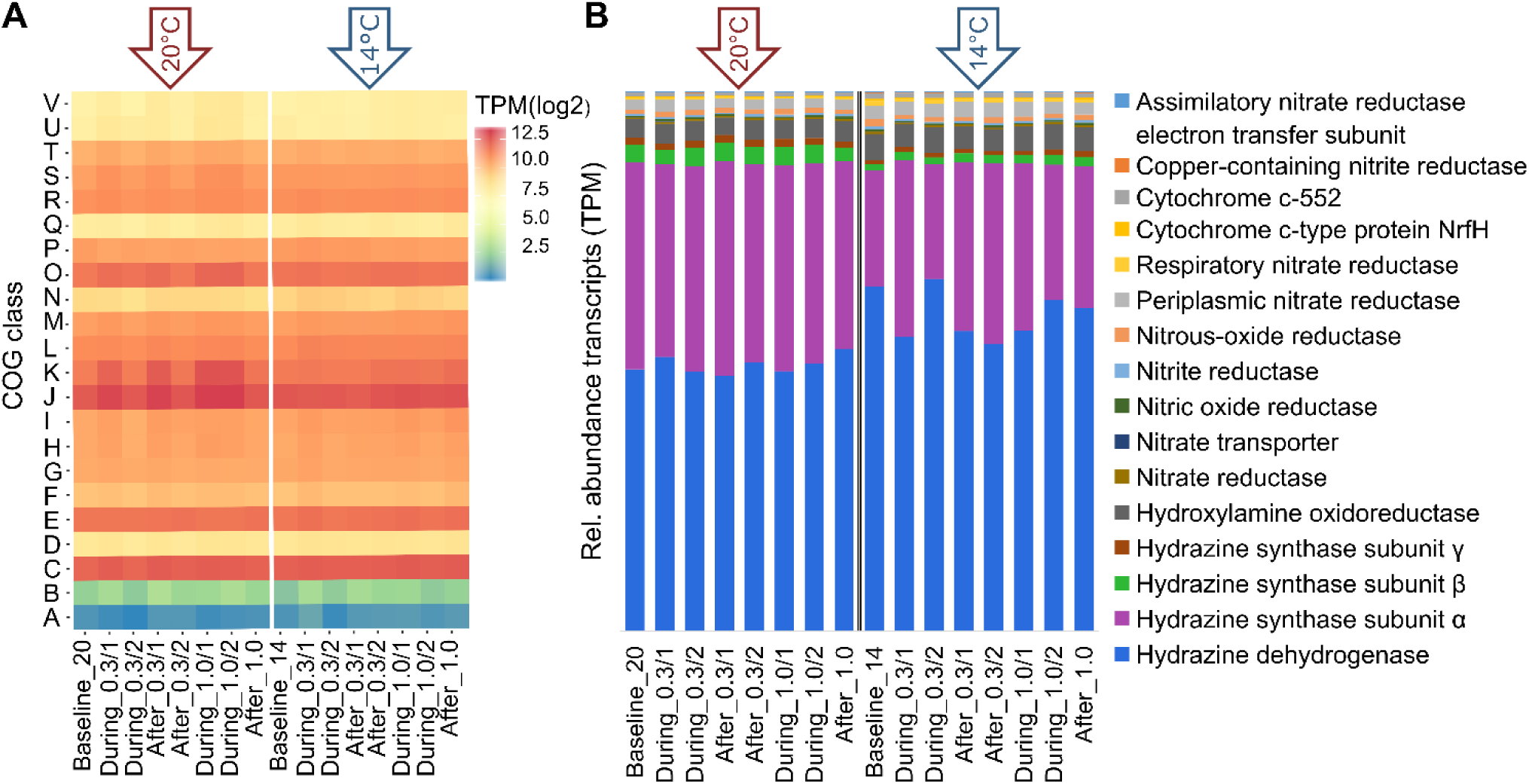
**A** Heatmap based on the cumulated transcripts per million (log2) from all genes falling into certain cluster of orthologues (COG) categories (y-axis). Time-points (x-axis) are based on the averaged reactors (n=3). **B** Relative abundance of transcripts in TPM of genes involved in the Nitrogen cycle. The Left part shows the 20 °C samples the right part the 14 °C experiment, as indicated by the arrows.

Transcript abundance confirmed the dominance of the anammox process already seen in the metagenome. Highest transcription was observed for genes involved in the anammox metabolism (Figure 5B). Notably, transcripts of hydrazine dehydrogenase and hydrazine synthase subunit alpha had up to two fold higher relative transcript abundance than other N cycle genes (Figure 5B). This is in line with previous findings on AMX genomes and corresponding transcript abundances^17, 31^.

Taking all N cycle genes into account, temperature again had an distinct effect on the transcriptional status of the community (see ordination in Supplementary Figure 7). While transcription of *hdh* genes was elevated at 14 °C, the transcription of *hzs* subunits α and β were significantly reduced (p < 0.05, Student’s t-test) (Figure 5B). In contrast to the community-level transcription trend, *hdh* displayed constant levels of transcription at 20°C, but more variations at 14°C. Similar to recent reports^17^, we also found moderate expression levels for partial or full denitrification. Genes encoding for denitrification and DNRA did not display significant differences between the temperature regimes.

### Transcriptional response of individual MAGs

Changes in the transcription at the level of functional genes in a microbial community as described above may arise from global changes of transcriptional activity in many different species as well as from specific transcriptional regulation within individual species. To investigate this, we combined MAG reconstruction with our transcriptomic approach.

Relative transcript abundances as a proxy for expression levels were calculated from the total filtered mRNA reads from 56 RNA samples, which were individually mapped against the recovered MAGs, normalized for respective genome length and sequencing depth and expressed as transcripts per million (Supplementary Table 2)^17, 58^. The two most abundant anammox MAGs (∼18% relative abundance) together were significantly (p < 0.05, Student’s t-test) responsible for the highest transcriptional activity during the 20 °C and 14 °C regime, respectively (Figure 6A). Less abundant *Candidatus* Brocadia MAGs (AMX3, AMX4, AMX5 and AMX6) demonstrated only moderate transcriptional activity. Temperature had the strongest effect on AMX5, which was more active at 14 °C while AMX2 appeared more active at 20 °C. To highlight the relationship between transcriptional activity and abundance we calculated the ratio between these two metrics for each anammox MAG. Ratios were comparable during the different temperature regimes except for AMX2, and AMX5. The high ratio of AMX2 can be attributed to a low abundance but high transcriptional activity.

**Figure 6.**
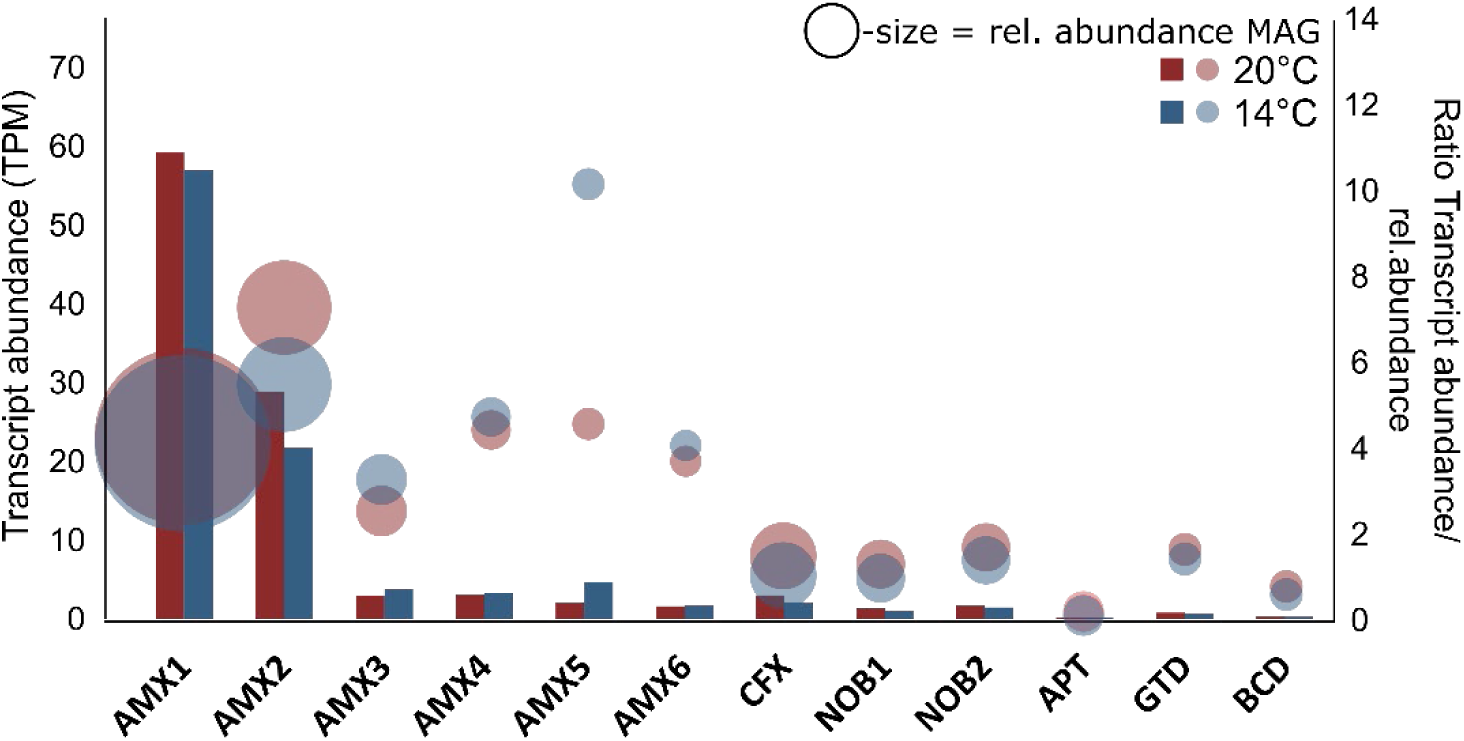
Barcharts representing the relative transcript abundance (in TPM) of MAGs recovered from the bioreactor. Bubbles represent the ratio between transcript abundance and corresponding relative abundance for each MAG. The second y-axis denotes the ratio levels. The size of bubbles correspond to the relative abundance of the MAGs. Supplementary Table 2 gives more details on mapping results.

After finding these differences in the transcriptional activity of the retrieved MAGs we compared transcriptional patterns within the functional classifications of COGs. Here, we summed up the TPM of all genes in each COG class for each MAG at each time-point. This dataset was investigated by ordination. For ease of interpretation, we split the nMDS (Jaccard dissimilarity) based on the temperature regimes (Figure 7). Transcriptional activity of COG categories shows MAGs to be clearly and consistently distinct, as evidenced by clusters within the nMDS space. We found that MAGs derived from the same functional group (e.g. AMX and NOB, which are in our case also from the same genus), were distinct from each other but clustered closely together while being more distinct from phylogenetically more distant MAGs (hull polygons in Figure 7). Furthermore, we found the degree of transcriptional change over time differed between individual MAGs (spread of points for each MAG in the nMDS space). Especially the spread between time points within the NOB cluster suggested dynamic changes in gene transcription independent of the temperature regime. Some MAGs displayed a clear temperature dependency in their transcriptional responses over the course of the experiment (Supplementary Figure 8). Specifically, within the AMX cluster, we found that temperature strongly affected the transcriptional profile of AMX3, AMX5 and AMX6, as well as the degree of transcriptional change (Figure 7, Supplementary Figure 8).

**Figure 7.**
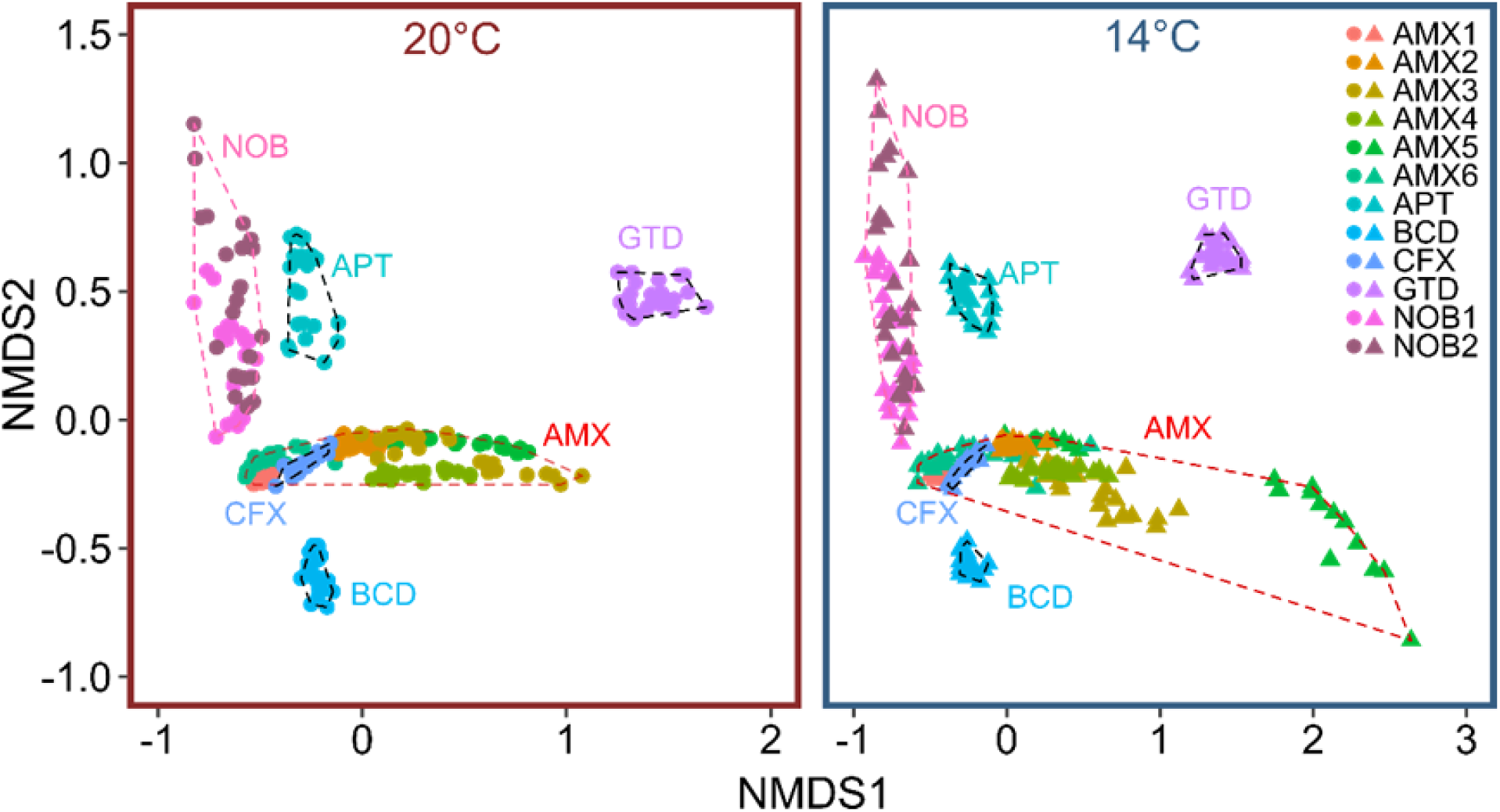
nMDS (based on Jaccard dissimilarity) based on the COG expression profile for each MAG. Colors denote different MAGs and dashed lines (ordihull function) highlight genus cluster. Each dot represents the COG profile of the corresponding MAG on a respective time-point. Left panel presents the 20 °C and right panel the 14 °C experiment. Stress value: 0.074

## Discussion

Using laboratory scale bioreactors and meta-omic approaches, we were able to investigate anammox process stability towards DO pulse disturbances and different temperature regimes on various ecosystem and biological levels.

While both temperature regimes started with similar N removal efficiencies, the lower temperature treatment had a significant longer-term negative effect on anammox performance. In agreement with previous findings on temperature induced performance loss in anammox reactors^23, 59, 60^, this highlights again that one of the main issues with the mainstream application of autotrophic N removal is the temperature sensitivity of anammox bacteria.

DO disturbances had a clear, but short-term reversible influence on the anammox process level. In strong contrast to our expectations, DO disturbances had a much more severe impact at 20 °C on the NH_4_^+^ removal rate; resulting in, depending on the DO concentration, partial to even full inhibition. During the 14 °C experiment, DO shocks did not lead to these drastic decreases in anammox activity. Previous studies that investigated the impact of DO on the anammox process also found immediate but reversible activity losses at higher temperatures (30 °C)^25, 26, 29^. This study, however, is for the first time comparing the DO response under different “cool” temperatures, which are more representative for mainstream conditions and to some extent mimic seasonal fluctuations in WWTP.

Looking beyond the black box of performance, our main aim was to understand the impact of disturbances during autotrophic N removal also at the level of transcription. In line with our findings on the process level, transcription was noticeably affected by disturbances at 20 °C, which is generally considered a more favorable temperature for anammox bacteria (Figure 4). This might indicate a higher capacity of the community to actively react to stress at this temperature and a restrained physiology and stunted stress response at 14 °C. The observed, elevated return times to the referential state after exposure to 1.0 mg L^-1^ h^-1^ DO under 14°C might also be an indication of temperature-restrained physiology.

Despite the pronounced temperature-dependent differences on the process and global transcriptional level, transcriptional responses to short-term DO disturbances were generally not strong nor did they lead to explanatory and /or indicative gene regulation cascades in the community as we had originally anticipated. However, categorizing genes into their respective COG class revealed a clear up-regulation of genes involved in the transcription, translation, replication and posttranslational modification associated with the DO disturbance under the 20 °C regime. Especially, genes involved in bacterial stress tolerance displayed the biggest fluctuations in expression due to DO disturbances. The up-regulation of heat-shock and cold-shock proteins allows for the maintenance of cellular processes during stress events and initiates a stress response cascade, which allows adaptation to harmful conditions^61, 62^. These stress response systems did not react during the 14 °C experiment. Paradoxically, given the short-term disturbances in our experiment, this lack of response may have contributed to a higher resistance of the process observed at 14 °C. In this experiment, no long-term activity loss was observed after the short return phase, indicating that even at less favorable temperatures the anammox consortia suffered no permanent damage from the oxygen stress levels induced here. However, we speculate that with repeated or stronger disturbances the lack of transcriptional response could result in reduced performance eventually. Transcript abundances were comparable during baseline conditions between (20 °C/14 °C), which suggests that the temperature drop or increase from the parent reactor (17 °C) to the experimental reactors did not impose an experimental background stress.

While NH_4_^+^ removal rate was clearly affected by DO disturbances, they did not suppress the transcription of any of the key N cycle genes within the system, independent of temperature. Similar transcriptional responses patterns towards oxygen perturbations have been observed for anammox bacteria from coastal systems^63^. Together with our results on the process level, this suggests inhibition took place on the post-transcriptional level. Further evidence for this hypothesis is the rapidity of the functional response which was independent of the temperature same for reaching a maximum inhibition - if inhibition would have happened on the transcriptional level, the reduction in process rates presumably would have been less abrupt, as residual proteins would have continued to function until internal enzyme levels became depleted. Similarly, it would have presumably taken much longer for the system to resume its initial NH_4_^+^ removal efficiency if protein synthesis would first have to be initiated. Experimental studies on denitrifying bacteria in pure culture observed a timeframe of 10-24 h to establish a fully active denitrification enzyme system after a shift from aerobic to anaerobic conditions ^64^. In our experiments, it took on average 37 min to return to referential state conditions. Based on our observations we speculate that during lower temperatures, not only the enzyme activity but also the enzyme inhibition is constrained which might have led to the less pronounced impact on the NH_4_^+^ removal rate during 14 °C. While the process response was thus not directly related to regulation of transcription, we nevertheless observed temperature-dependent differences in the transcription of key enzymes involved in the anammox process. We argue that the significantly higher expression of the *hdh* gene and lower expression of the *hzs* gene during the 14 °C experiment is probably an adaptation mechanism to lower temperatures. Hydrazine is toxic to anammox microorganisms if accumulated for long periods inside cells^65^. At colder temperatures, the metabolic machinery of anammox bacteria slows down, and this may affect various steps of the process differently. To avoid hydrazine accumulation, the anammox cells appear to use transcriptional regulation to increase hydrazine dehydrogenase enzyme production and decrease hydrazine synthase enzyme production to ensure that all toxic hydrazine is converted efficiently to inert N_2_. Reduced hydrazine synthase expression might have also contributed to the slow but steady performance loss during the 14 °C treatment.

We are aware that MAGs are computationally constructed entities and that caution should be taken in interpreting them as representative biological species. However, in order to investigate stress response of individual members from complex microbial communities we consider this approach will offer new scientific insights that are also relevant for the applied science field. It offers new opportunities to further our understanding how species-driven responses lead to process stability or failure in engineered ecosystems.

Using this novel approach, we found that with only 19% relative abundance, anammox MAGs (AMX1-6; all *Candidatus* Brocadia) were nevertheless displaying highest transcript abundance in the system and drive the majority of the community metabolism, independent of the temperature regime. Interestingly, this dominance in community transcript abundance was also reported for anammox bacteria in a laboratory-scale reactor, operated under sidestream conditions (30 °C), but here they accounted for ∼ 65% of the overall community^17^. These findings again support the notion that anammox bacteria, despite lower abundance and unfavourable temperatures, can achieve efficient N removal under mainstream-like conditions^66^.

The presence of numerous *hao*-like genes and lack of *nir* genes in the retrieved anammox MAGs suggests that they pursue the recently proposed hydroxylamine-dependent anammox mechanism^67^. Here, NO_2_^-^ is reduced to hydroxylamine, which is then, together with NH_4_^+^, converted to hydrazine. The taxonomic classification of the MAGs as members of the *Candidatus Brocadia* spp. lineage supports this notion, as none of the previously studied representatives encode any known nitrite reductases^68^. Furthermore, we found that all anammox MAGs contain the nitric-oxide reductase gene (*nor*), which produces the homonymic enzyme, which reduces NO to N_2_O and potentially detoxifies the system^69^. We did not observe increased gene activity of *nor* on the community level, which might be explained by the non-inhibiting but rather beneficial effect of NO on anammox bacteria^70^.

The other retrieved MAGs together with the anammox MAGs shaped the backbone of a nearly closed N loop and could enhance overall N removal in the bioreactor^18^. Two-stage denitrification paired with anammox is a promising alternative to maximize biogas production^71^. However, it could also facilitate competition for denitrification-intermediates between anammox bacteria and other N key-players in the system^17^.

Focussing on the stress response of individual MAGs revealed characteristic temporal COG transcription profiles for each MAG but also similarities between closest phylogenetic relatives. Stress response on the transcriptional level can be distinct within genera but seems to be significantly different between phyla. While the two most dominant anammox MAGs displayed overall stable transcript abundance, transcriptional change was much larger during the 14 °C experiment for less abundant anammox MAGs. The biofilm carriers used in our experiment were exposed to mainstream ambient temperatures and seasonal temperature fluctuations for a period of two years. The most abundant species of the AMX consortia presumably have adaptations that allow them to thrive under these unstable conditions^72^. The low abundant AMX MAGs on the other hand are generally stressed due to constant competition for available resources with the abundant AMX fractions^73^. Thus, additional stress might have imposed the elevated stress responses of these MAGs.

## Summary and outlook

Further studies of more severe stress and even actual system failures are needed to better understand the consequences of disturbance on diverse microbial community and transcriptional levels within these engineered ecosystems. Here, temperature was shown to change microbial community status in complex ways, but AMX biofilms proved resilient against short O_2_ disturbances independent of temperature.

Long-term exposure to colder temperatures or prolonged DO disturbances might lead to completely different dynamics within the community but also on the individual level. We believe that our findings on transcriptional stress response advances our insight on the links between microbial community stress response, individual stress response and process level failure. Furthermore, it emphasizes the value of molecular techniques paired with cutting-edge bioinformatics to understand individual stress response of key players in engineered ecosystems.

## Acknowledgments

This work was supported by funding from the Swiss national science foundation Synergia project ISOMOL: CRSII5_170876. The authors would like to thank Moritz Lehmann (University of Basel) and Joachim Mohn (EMPA) for the helpful scientific discussions during the whole period of this study.

## Author contributions

R.N, D.H, A.J and H.B designed the study. R.N and D.H performed the experiments. R.N performed the sampling, sequencing, data analysis and wrote the first draft of the manuscript. A.P contributed the binning to this study. R.N and H.B wrote the manuscript with critical and helpful reviews from D.H, J.W, A.P, P.M and B.S.

## Additional information

### Competing interests

The authors declare no competing financial interest

