## Supplementary Information for "Temperature modulates stress response in anammox reactors"

**This file includes:**

**Supplementary figures 1-8**

**Supplementary tables 1-2**

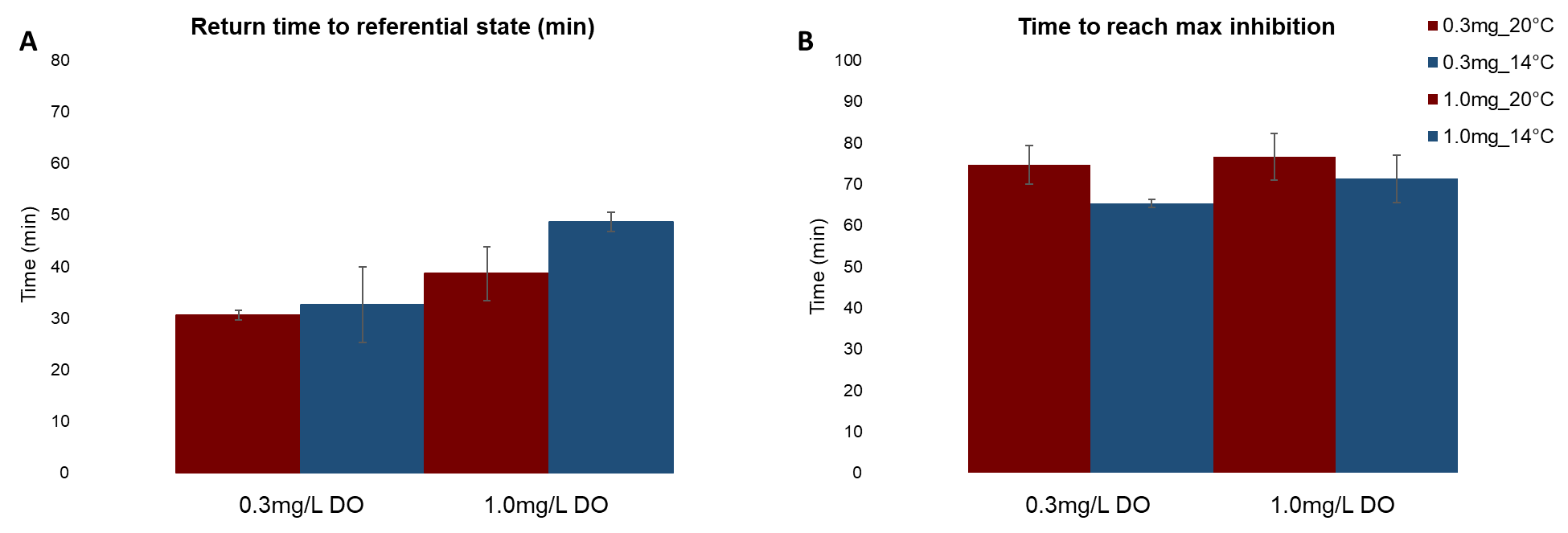

Supplementary Figure 1 **A** Return time to baseline performance levels (NH_4_^+^ removal rates) in minutes after dissolved oxygen perturbations under different temperature regimes (20 °C, red; 14 °C, blue). Left bars denote 0.3 mg L^-1^ DO stress response while right bars reflect the 1.0 mg L^-1^ DO stress response. **B** Time in minutes to reach maximum impact of the applied DO disturbance. Triplicate reactors were averaged for this graph.

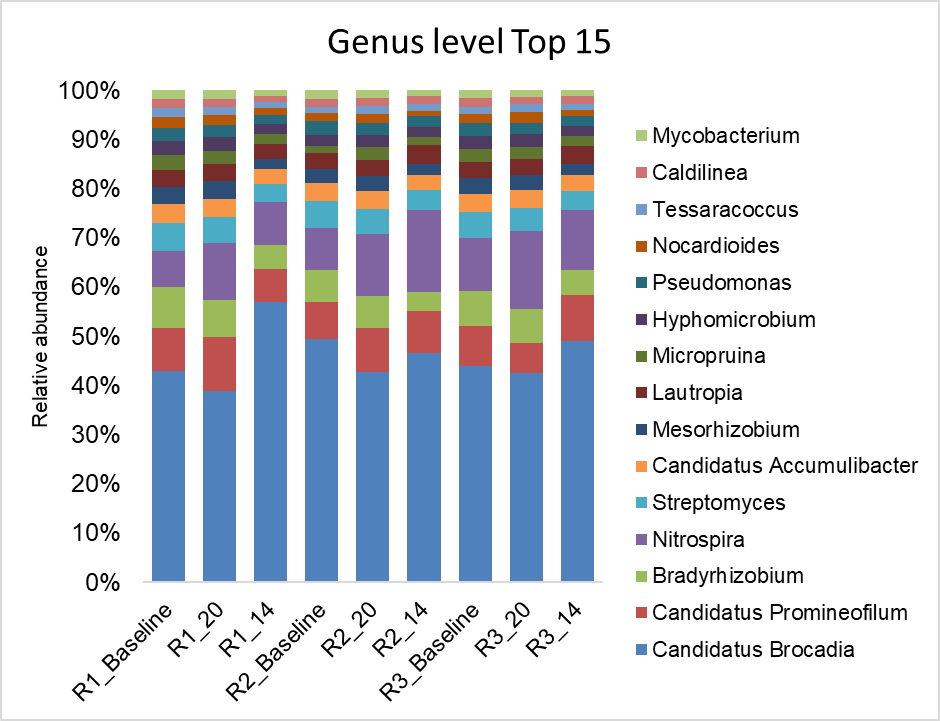

Supplementary Figure 2 Relative abundance of the top 15 bacterial genera of the biofilm carrier community derived from metagenomic sequencing. Labels denote Reactor (R1, R2, R3) and time of sampling, _**Baseline** (Pooled from 20 °C and 14 °C experiment), **_20** (after the 20 °C experiment), **_14** (after the 14 °C experiment). Colors represent the different bacterial genera.

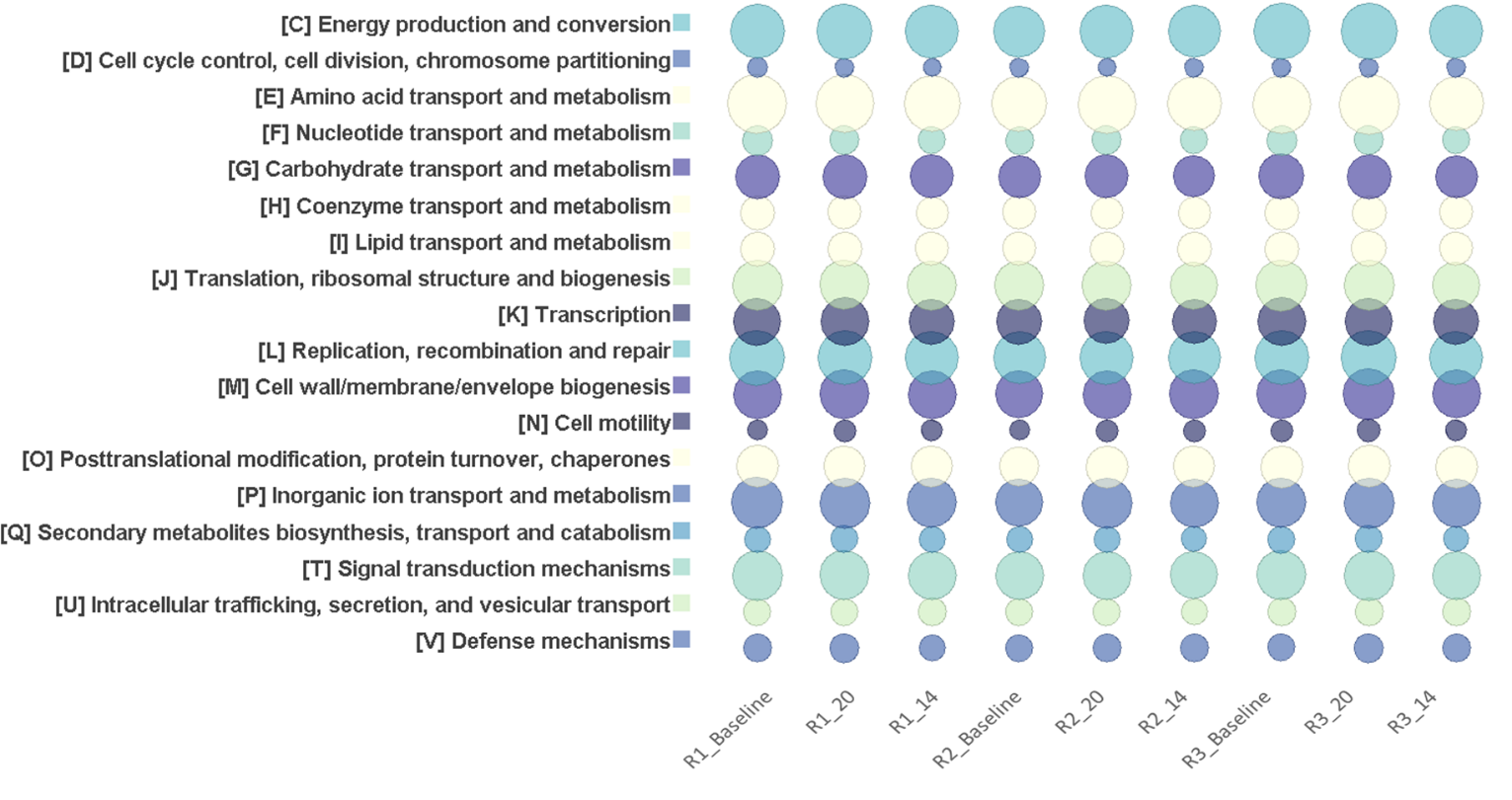

Supplementary Figure 4 Gene abundances categorized into **Seed subsystems**. Labels denote Reactor (R1, R2, R3) and time of sampling, _**Baseline** (Pooled from 20 °C and 14 °C experiment), **_20** (after the 20 °C experiment), **_14** (after the 14 °C experiment). Size of bubbles correspond to the relative abundance of the system.

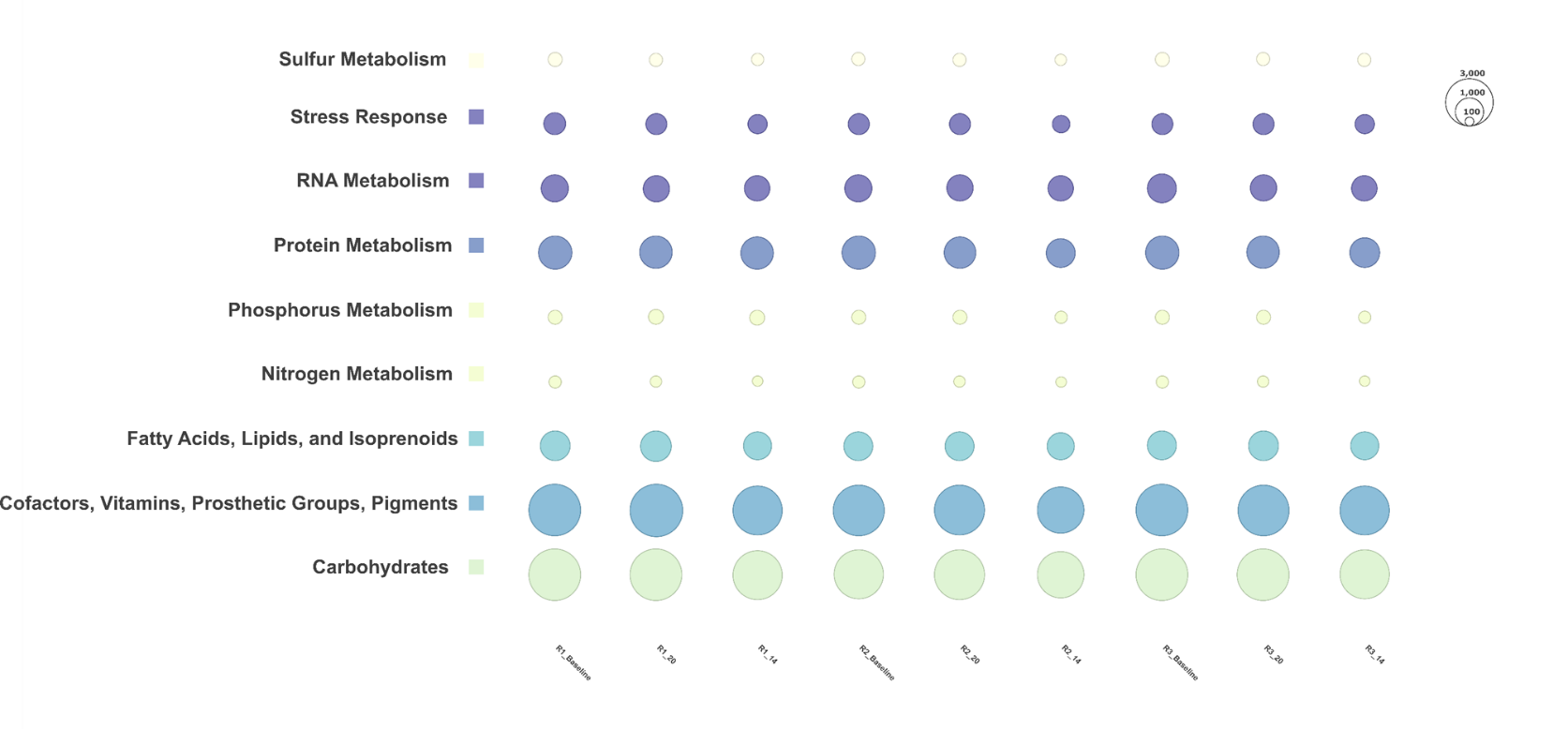

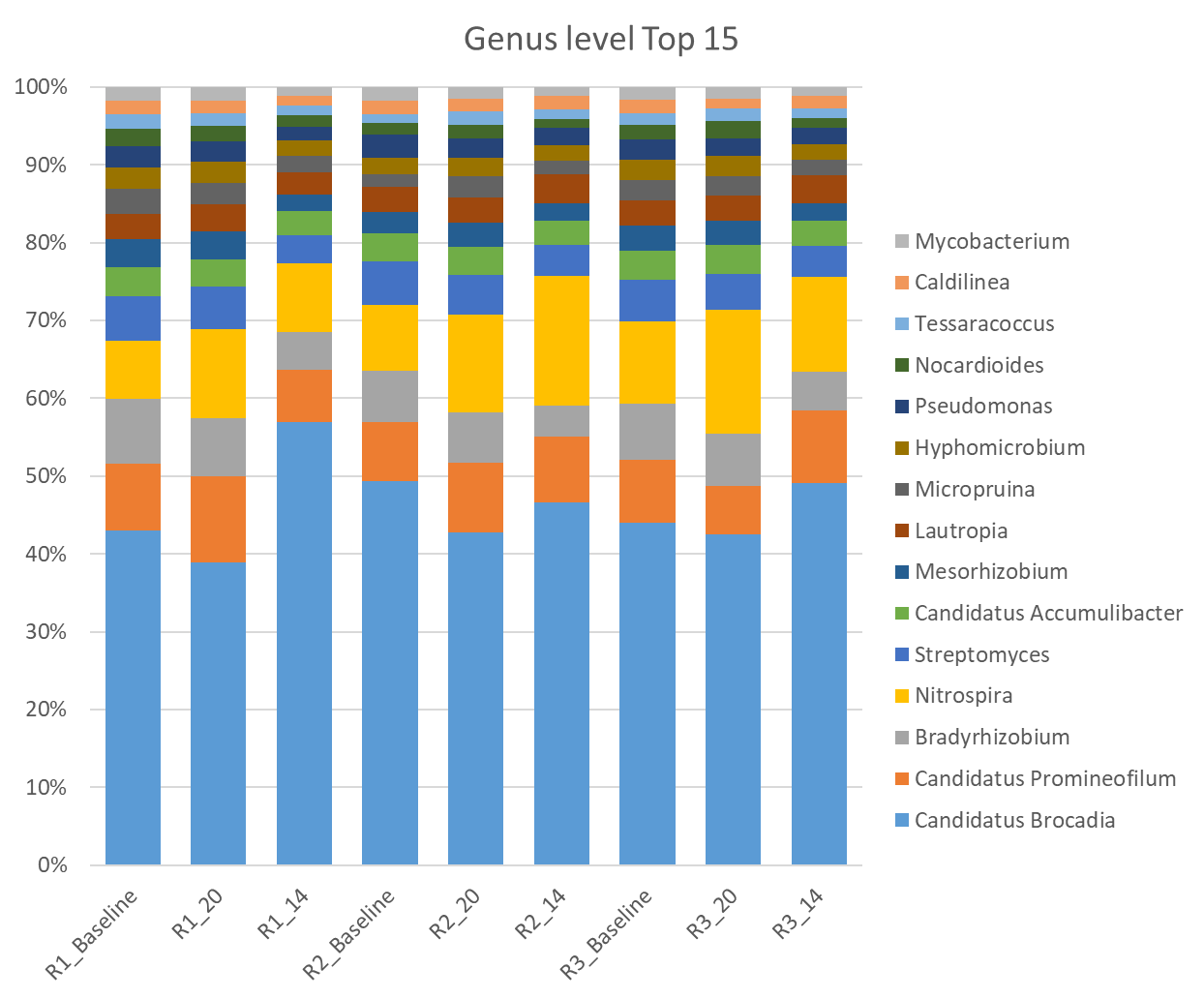

Supplementary Figure 3 Gene abundances categorized into **C**luster of **O**rthologues **G**roups (COG). Labels denote Reactor (R1, R2, R3) and time of sampling, _**Baseline** (Pooled from 20 °C and 14 °C experiment), **_20** (after the 20 °C experiment), **_14** (after the 14 °C experiment). Size of bubbles correspond to the relative abundance of the group.

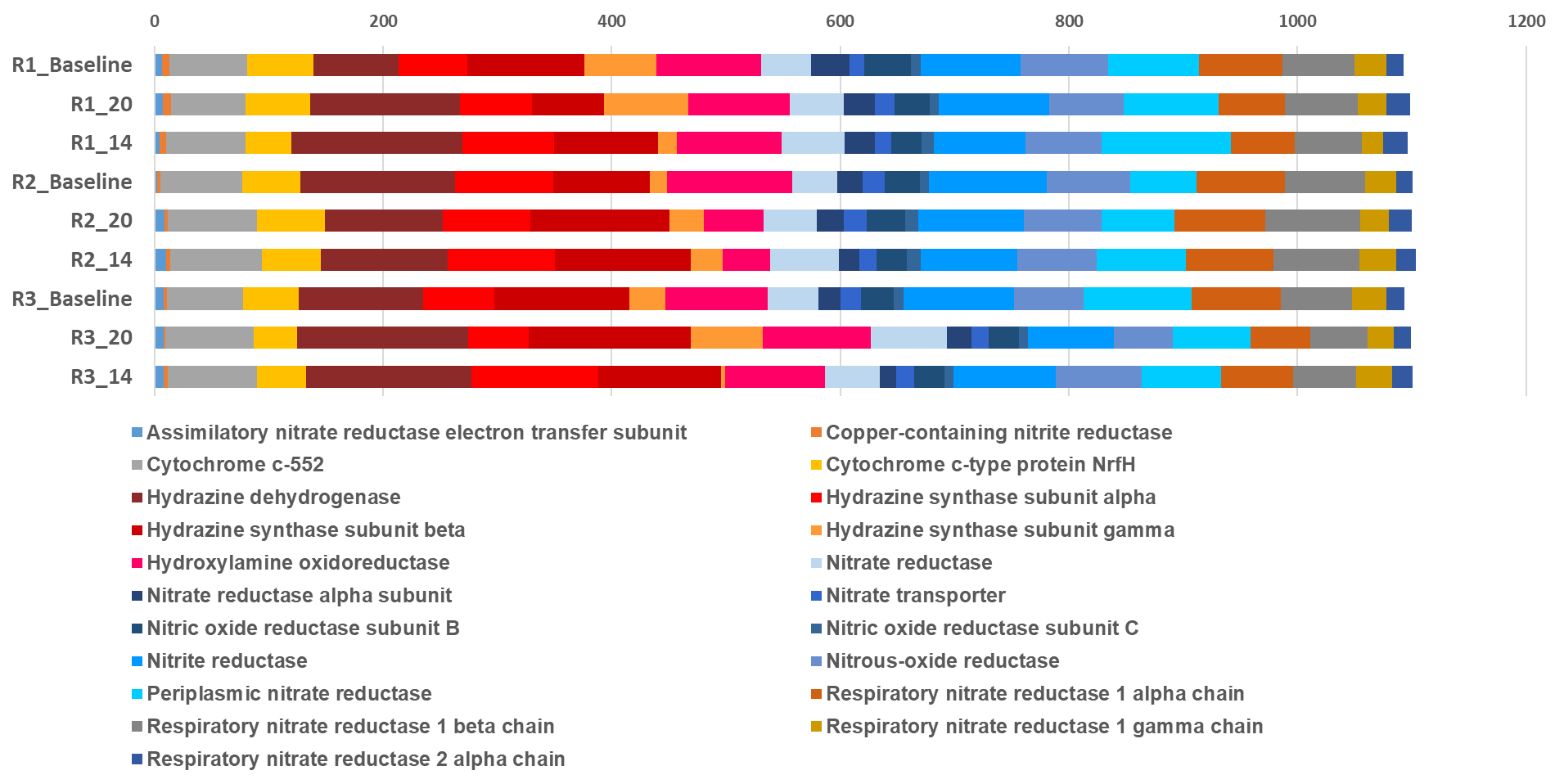

Supplementary Figure 5 Abundances of the most prominent genes from the nitrogen cycle expressed as genes per million. Reddish colours correspond to genes involved in the anammox cycle, blueish to the denitrification pathway. Labels denote Reactor (R1, R2, R3) and time of sampling, _**Baseline** (Pooled from 20 °C and 14 °C experiment), **_20** (after the 20 °C experiment), **_14** (after the 14 °C experiment).

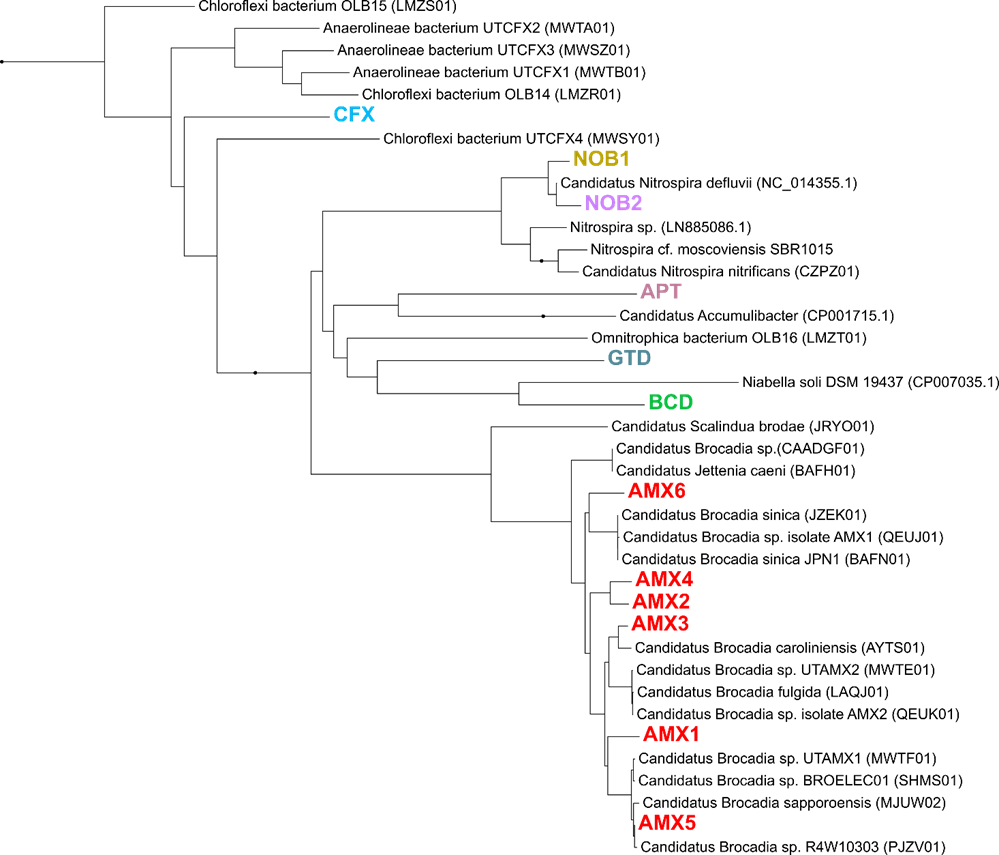

Supplementary Figure 6 Phylogenetic tree of all recovered draft genomes from the anammox bioreactor. Tree includes MAGs recovered from this study (different colours) and closely related genomes downloaded from the NCBI genome repository. GenBank accession numbers for each genome are provided in parentheses. The tree was constructed using RAxML based on a set of 37 concatenated universal single-copy marker genes.

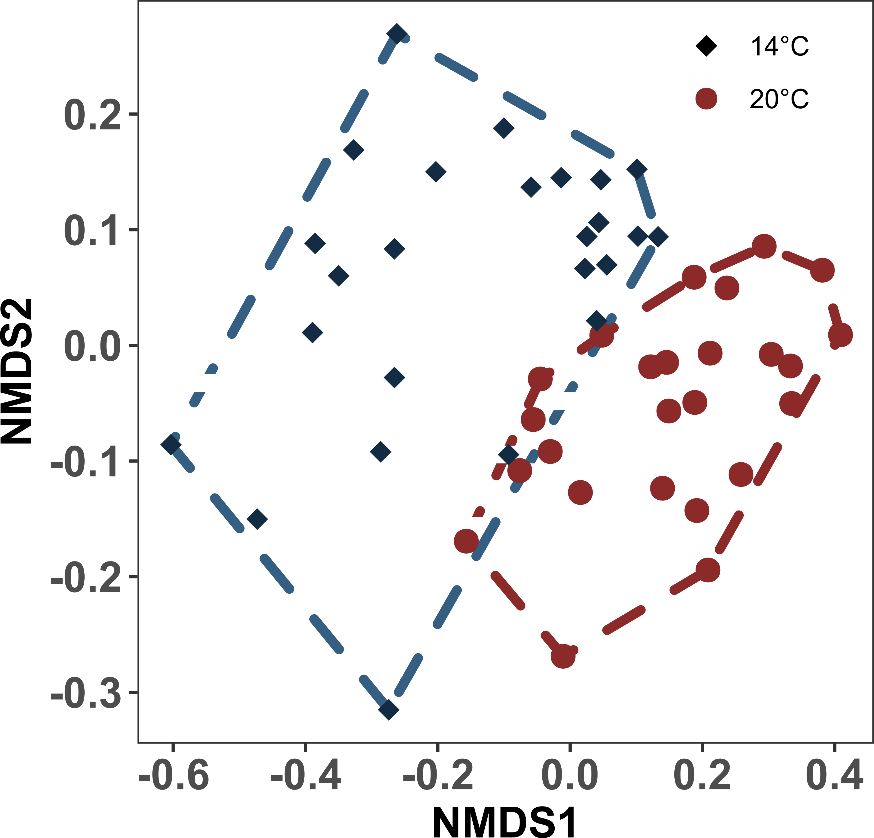

Supplementary Figure 7 nMDS based on Jaccard dissimilarity depicting all genes involved in the Nitrogen cycle. Each dot represents a timepoint. The colour denotes the temperature regime. Hulls highlight also the temperature regime. Stress: 0.115

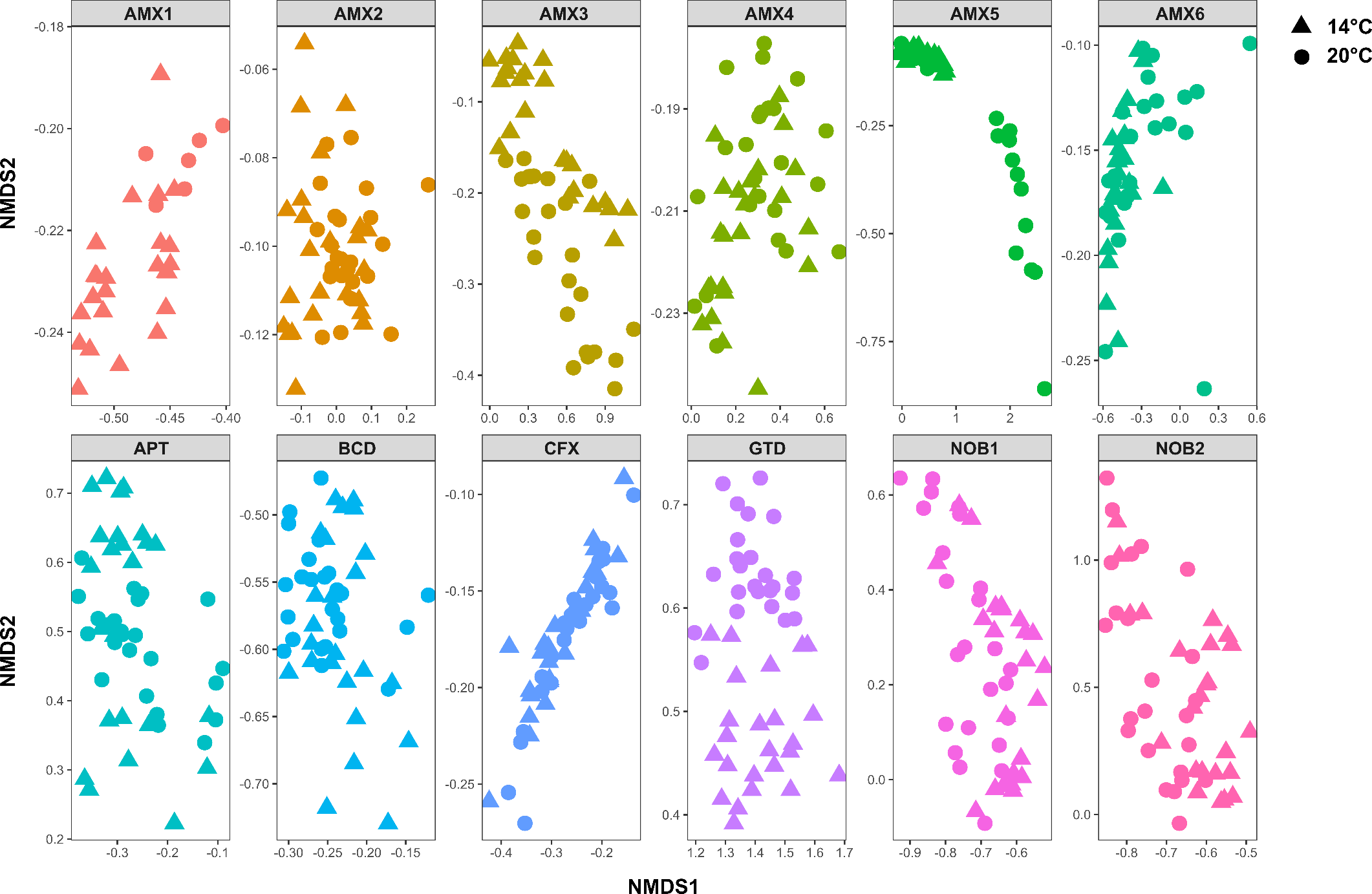

Supplementary Figure 8 nMDS (Jaccard dissimilarity) based on the COG expression profile for each MAG. Colors denote different MAGs shapes correspond to the temperatures. Each dot represents the COG profile of the corresponding MAG on a respective time-point. Symbols denote the temperature (dot 20 °C; triangle 14 °C). Stress: 0.074

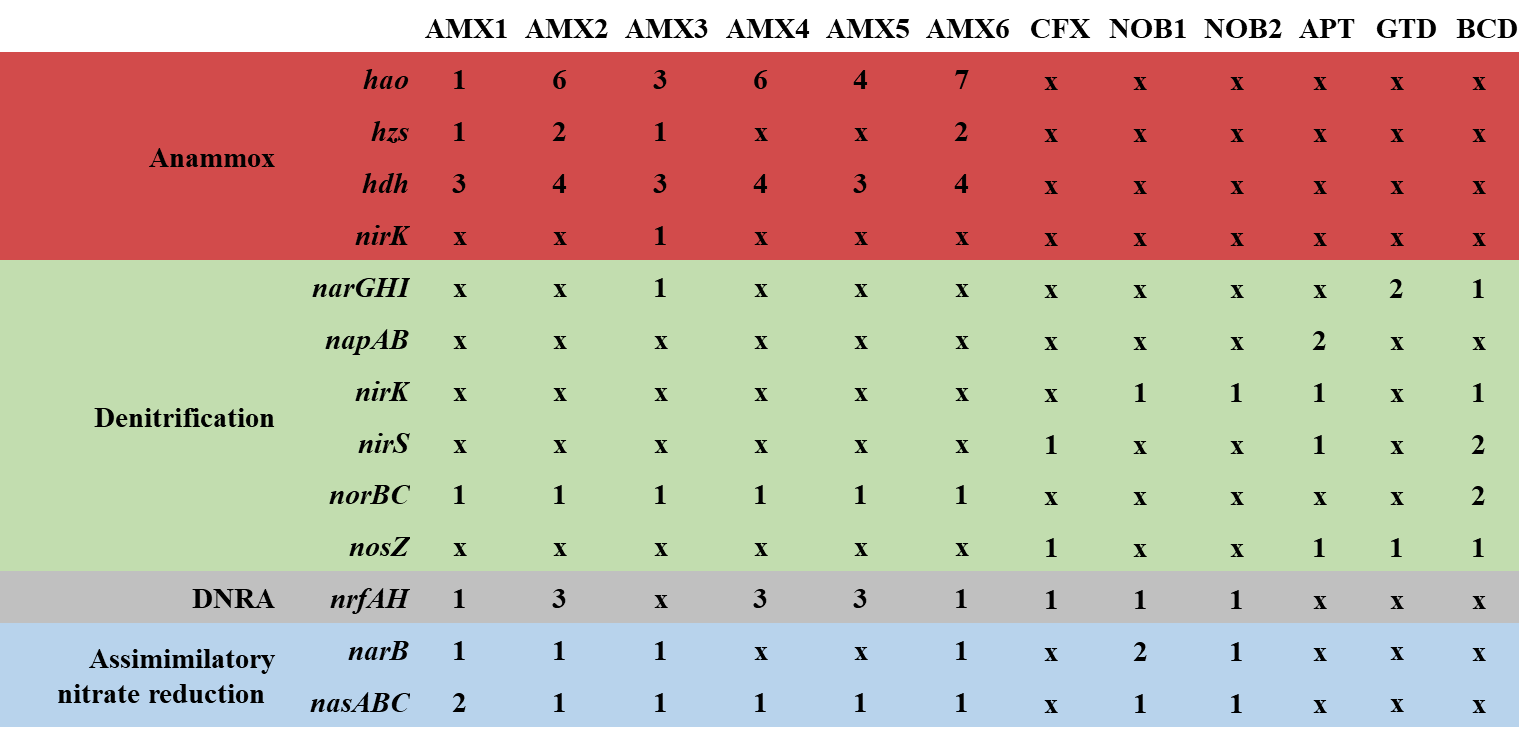

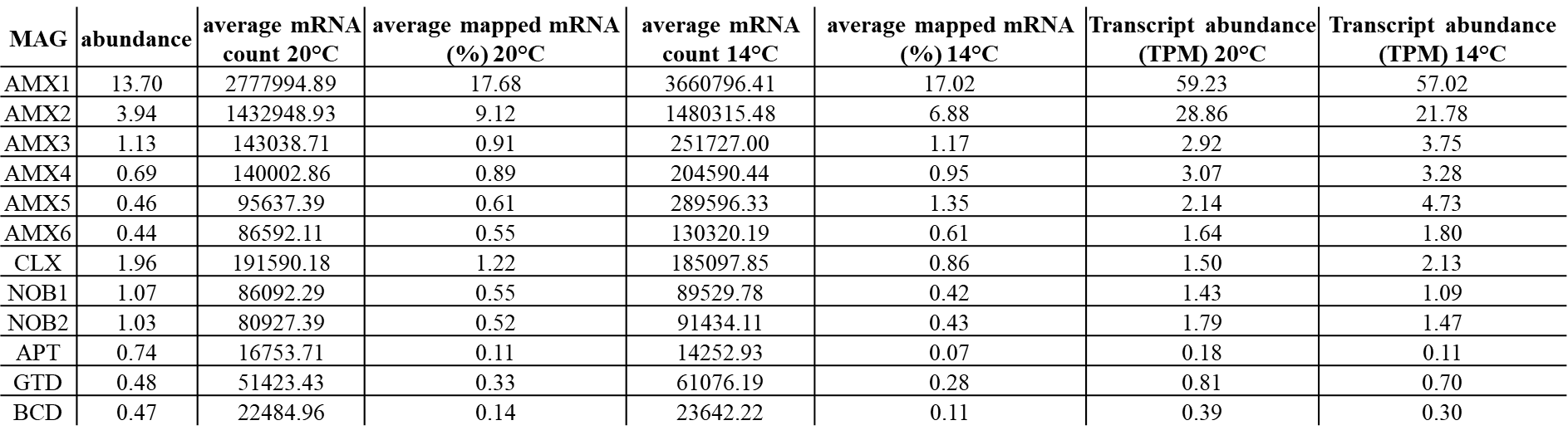

Supplementary Table 2 Relative Abundance of MAGs and gene expression estimates, based on transcripts per million values of transcripts that mapped to each MAG

Supplementary Table 1 Nitrogen cycle genes presence/absence in metagenome assembled genomes. Numbers correspond to the amount of gene copies found within the respective genome.
